# DNA:RNA hybrid mutational landscapes reveal a common route to genetic instability in evolving genomes

**DOI:** 10.64898/2026.09.25.754379

**Authors:** Myriam Zheng, Fanny Pouyet, Samuel O’Donnell, Nicolas Agier, Raphaël M. Mangione, Louis Ollivier, Christie Ouaddi, Umberto Aiello, Coralie Goncalves, Ophélie Lautier, Anna Babour, Domenico Libri, Gilles Fischer, Benoit Palancade

## Abstract

Pervasive DNA:RNA hybrids are recognized as pathological sources of genome instability, yet how their genotoxicity shapes mutational landscapes and genome evolution remains poorly understood. Here, we define the mutational footprint of hybrids by integrating genome-wide mapping, long-term mutation accumulation experiments, and analyses of genetic diversity across thousands of natural yeast genomes. We uncover signatures for embedded ribonucleotides and genic R-loops, together with a composite mutational pattern shared across natural populations and experimental evolution. Using reporters designed to dissect the mechanisms underlying genetic alterations, we find that they arise primarily from targeting by MutLγ-dependent mismatch repair followed by Rad52-mediated recombination. Although hybrids form dynamically across the genome, their genotoxicity is strongly skewed toward repetitive regions susceptible to these error-prone pathways, including microsatellites and retroelements. Our findings reveal a common mechanism of hybrid-associated mutagenesis that can shape genetic variation in evolving genomes.

## Introduction

Mutations are the driving force behind the evolution of genomes and the development of pathologies, and often arise from the error-prone processing of DNA lesions during replication or repair. Among the intrinsic sources of mutations and genetic instability are DNA:RNA hybrids, which range from single ribonucleotide insertions incorporated during replication^1^, to longer hybrids formed during transcription as three-stranded R-loops, in which hybridization of nascent RNA to its DNA template displaces the non-template DNA strand^2,3^. Advances in genome-wide mapping techniques^4–6^ have revealed that these structures are widespread across model organisms and impact a substantial fraction of the genetic material^7–10^. Accordingly, their homeostasis is maintained by conserved pathways that either remove existing hybrids, notably RNases H and helicases, or prevent their formation through RNA-binding protein complexes such as the THO/TREX complex and the spliceosome^11–13^. Defects in DNA:RNA hybrid regulators have been linked to pathologies, including cancers and neurodegenerative disorders^14,15^, but their contribution to disease remains difficult to establish because these factors often exert pleiotropic effects, including widespread transcriptome alterations^16–18^.

Both excess ribonucleotides and DNA:RNA hybrids are generally recognized as a source of genetic instability, either because they are targeted by DNA metabolism enzymes^19–25^ or because they trigger replication stress^26–28^. While extreme hybrid accumulation is lethal^27–29^, changes in hybrid levels have been associated with different types of genetic alterations, essentially through the use of reporter systems or ensemble assays. Both types of hybrids can promote mutations, *i.e.* single nucleotide variants (SNVs) and insertions/deletions (INDELs)^19,20,30,31^, and are also recombinogenic, triggering double-strand breaks (DSBs) and, ultimately, gross chromosomal rearrangements (GCRs), as revealed by dedicated reporters^32–34^. In yeast, DNA:RNA hybrid-dependent rearrangements have been shown to involve different homologous recombination pathways depending on the reporter system used, including single-strand annealing (SSA)^35^ and break-induced replication (BIR)^36^. Of note, while synthetic reporter systems are instrumental to describe the mechanisms by which these structures drive genetic instability, they have limitations to evaluate their impact on natural, unstressed genomes. Indeed, their sequences, organization and expression programs may have coevolved to minimize the deleterious effects of DNA:RNA hybrids, as recently suggested in bacteria^37^. A substantial body of evidence indeed supports that not all DNA:RNA hybrids are equally genotoxic, with fewer than 20% of hybrids being associated with DNA damage features in both yeast and human cells^29,38^. Further complicating the assessment of their genotoxicity, hybrids have also been shown to accumulate subsequently to replicative stress or DNA damage^39–43^, and, in some cases, to benefit genome maintenance^44^. Understanding which of these hybrid subtypes contribute to genome instability and genetic variation requires the systematic analysis of the mutations arising from hybrid accumulation, as achieved previously under perturbation of several other DNA metabolic processes^45,46^. For this purpose, we combined the systematic mapping of DNA:RNA hybrids, and the profiling of their associated mutational signatures, exploiting both the genetic diversity of natural populations, and the experimentally-driven accumulation of mutations in laboratory strains.

## Results

### A genic hybrid-associated mutational signature in natural yeast genomes

To determine the association of DNA:RNA hybrids with genetic alterations, we first examined the distribution of hybrids within the yeast genome, by performing calibrated, strand-specific DNA:RNA hybrid immunoprecipitation followed by sequencing (qDRIP^47^) with the hybrid-specific S9.6 monoclonal antibody, using cells either wild type (*wt*) or inactivated for both RNase H1 and RNase H2 (*rnh1Δ rnh201Δ*), as we previously implemented^41^. Strand-specific signals were highly correlated between biological replicates, sensitive to *in vitro* RNase H treatment, reflecting their *bona fide* DNA:RNA hybrid nature, and consistent with available hybrid maps based on the utilization of the S9.6 antibody^41,48^ (**Ext. Data Fig. 1a-c**). In *wt* cells, DNA:RNA hybrid peaks were restricted to template strands and delineated by transcription start sites (TSS) and transcription end sites (TES), consistent with co-transcriptional hybrid formation, as evident in typical snapshots (**Fig. 1a**) and metagene analyses (**Fig. 1b**, **Ext. Data Fig. 1b**). The inactivation of RNases H led to the widespread accumulation of hybrids over pre-existing hybrid-forming regions (**Fig 1a-b**), demonstrating that our mapping approach identifies transcription-borne, dynamic hybrids that are normally resolved by RNases H in *wt* cells. Importantly, loss of function of RNases H barely impacted the transcriptome in our genetic background, with few loci exhibiting significant changes by RNA-seq (**Ext. Data Fig. 1d**). Genes grouped according to hybrid enrichment showed similar, limited changes in gene expression (**Ext. Data Fig. 1e**), indicating that the increased hybrid signals scored in the *rnh1Δ rnh201Δ* mutant do not simply reflect transcriptional upregulation.

**Figure 1.**
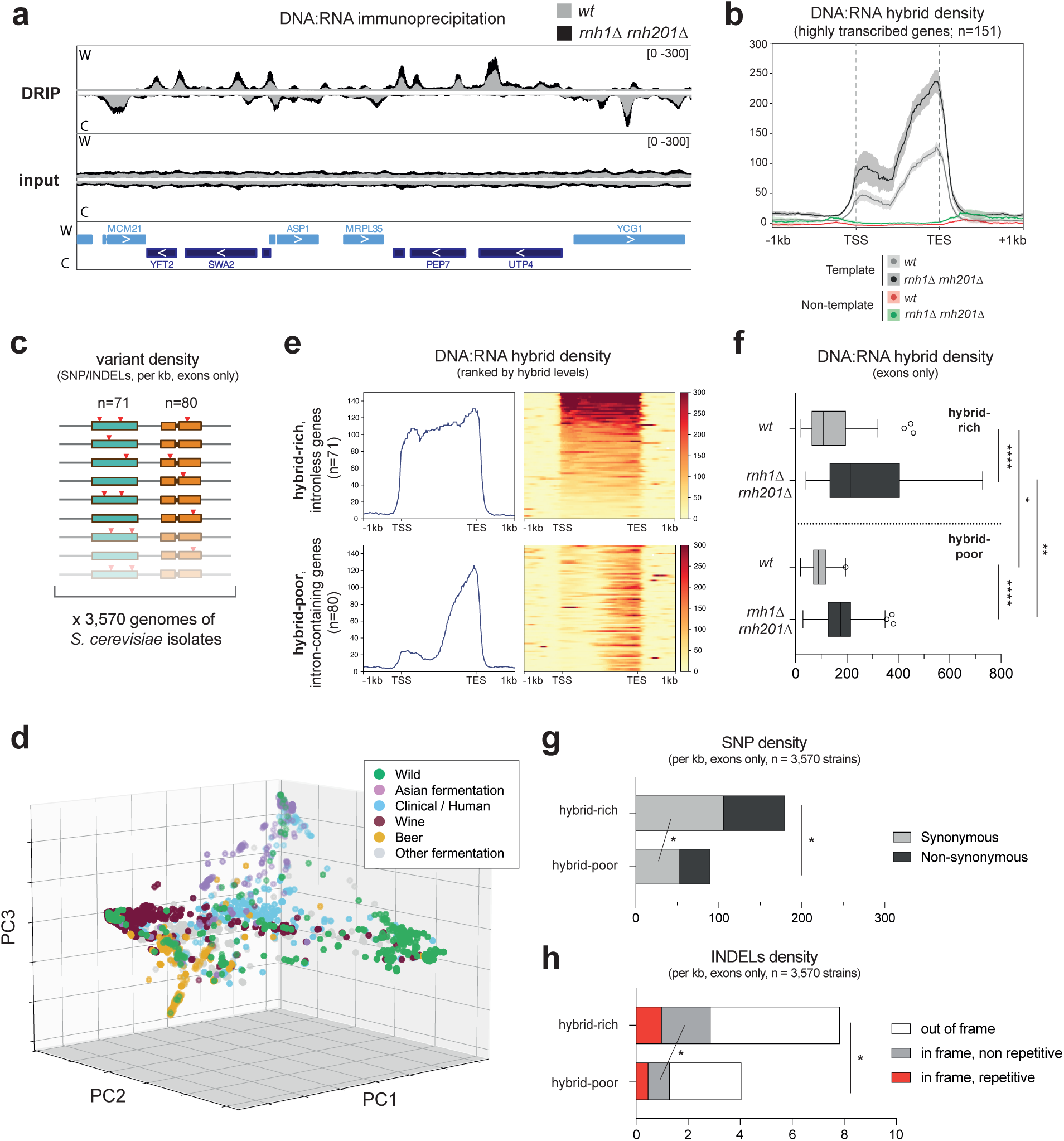
A hybrid-associated mutational signature in yeast populations. **a,** Integrative Genomics Viewer (IGV) representative screenshot of strand-specific DRIP-seq coverage in cells either *wt* (grey) or *rnh1Δ rnh201Δ* (black). Signals from Watson (W) or Crick (C) strand DNA from the region 1103117-1120436 in chromosome IV are represented. Average signals for DRIP replicates (*DRIP*) and total genomic DNA (*input*) are represented. **b**, Metagene analysis of DNA:RNA hybrid levels (DRIP signals in immunoprecipitates) at highly-transcribed genes (n=151) aligned at their Transcription Start Site (TSS) and Transcription End Site (TES). Averaged signals on template and non-template strand are represented for both *wt* and *rnh1Δ rnh201Δ* DRIP experiments. **c**, Principle of the population analysis. Variant density was assessed in gene sets comprising either hybrid-rich genes (blue; n=71) or hybrid-poor genes (orange; n=80) across 3,570 whole-genome sequences from distinct *S. cerevisiae* isolates. **d,** Principal component analysis performed on a list of 25,228 SNPs summarizing genome-wide patterns of genetic variation among the 3,570 strains. The first three principal components, which capture the largest fractions of the total genetic variance, are shown (PC1: 21.6%, PC2: 7.9%, PC3: 4.3%). Each point represents a strain colored according to its ecological origin. **e,** Metagene and heatmap analysis of DNA:RNA hybrid levels (DRIP signals in immunoprecipitates) at both hybrid-rich and hybrid-poor gene sets, aligned at their TSS and TES. Averaged signals on template strands are represented for *wt* DRIP experiments. **f,** Quantification of strand-specific DNA:RNA hybrid levels densities over transcribed regions (TSS-TES, excluding intronic sequences) in the hybrid-rich group (*top panel*) and the hybrid-poor group (*bottom panel*) in *wt* and *rnh1Δ rnh201Δ* DRIP experiments. **g**, SNPs densities (per kb, excluding intronic sequences) in the hybrid-rich and hybrid-poor gene sets. Non-synonymous and synonymous substitutions are represented. **h**, INDELs densities (per kb, excluding intronic sequences) in the hybrid-rich and hybrid-poor gene sets. INDELs are classified as out of frame, in frame (non repetitive) and in frame (repetitive). * p<0.05; ** p<0.01; **** p<0.0001 (**f**, Mann-Whitney-Wilcoxon rank sum test; **g-h**, Fisher exact test).

To analyze the correlation between hybrid signals and sequence polymorphisms within yeast populations, we compared variant densities between a hybrid-forming gene group (**Fig. 1c**, in blue) and a control, hybrid-poor gene group (in orange) across the genomes of 3,570 *S. cerevisiae* covering the geographical, ecological and genetic diversity of the species (**Fig. 1d**). The hybrid-rich gene group encompasses loci (n=71) which are both highly-expressed and intronless, two primary determinants of hybrid formation in the yeast genome, and which account for around 25% of the transcriptional output of *wt* cells^11,48^. As a control, we used an equally-sized group (n=80) of hybrid-poor, intron-containing genes that have similar transcription rates and base content (**Ext. Data Fig. 1f**), yet reduced hybrid accumulation and RNase H binding^11,49^. Analysis of our DRIP-seq datasets confirmed that hybrid-prone intronless genes displayed higher hybrid density as compared to their intron-containing counterparts in *wt* cells **(Fig. 1e)**, a difference which was further enhanced upon inactivation of RNases H **(Fig. 1f)**. To exclude transcriptional - and therefore hybrid - differences between isolates for the genes analyzed, we examined mRNAs expression levels for both gene groups across available transcriptomes^50^ for a subset of strains (n=969). Only marginal and comparable variations in mRNA levels were observed between strains for the two groups (**Ext. Data Fig. 1g**). Comparing these two gene sets therefore enables assessment of the consequences of hybrid differences that are consistently maintained across strains.

Variant analyses within exonic regions in these two groups then identified a total of 12,793 single nucleotide polymorphisms (SNPs) and 560 INDELs, both of which were strikingly more frequent in the hybrid-prone gene set than in the matched control group **(Fig. 1g-h)**. To control for the possible confounding effect of selection on these differences, we next focused on the more neutral synonymous SNPs and in-frame INDELs. Importantly, the same difference between the two groups remained detectable (**Fig. 1g-h**, *grey bars*), supporting the idea that hybrid-prone genes undergo enhanced mutagenesis as compared to their hybrid-poor counterparts. Of note, a substantial subset of the in-frame INDELs impacted repetitive sequences (**Fig. 1h**, *red bars*), corresponding to expansions or deletions within homopolymers, short trinucleotide repeats or other genic repeats in coding regions (**Ext. Data Fig. 1h**). Altogether, our data thus support that natural yeast genomes display signatures of hybrid-associated mutagenesis.

### The mutational signature of DNA:RNA hybrids appears in laboratory evolution experiments

To directly assess the impact of hybrids on mutational landscapes, we analyzed the genetic alterations spontaneously arising in the hybrid-forming *rnh1Δ rnh201Δ* mutant using long-term mutation accumulation (MA) lines (**Fig. 2a**). For both *wt* and mutant haploids, at least fifteen G_0_ colonies were streaked onto non-selective plates to isolate colony-forming units (G_1_), and further propagated through 25 successive bottleneck passages, with each passage representing ∼23 divisions (**Ext. Data Fig. 2a**), thus yielding a total of ∼600 generations per line and ∼9,000 generations per genotype. Whole genomes of G_25_ isolates were sequenced at high coverage (>250X) using short-read Illumina sequencing and compared to their G_0_ counterpart to detect all the mutational events appeared during the course of the experiment, including base substitutions (single nucleotide variants, SNVs), INDELs (<100bp) and other genetic alterations such as structural variants (SV, *e.g.* deletions, translocations), changes in ploidy and chromosome numbers (including aneuploidies), and mitochondrial DNA loss (**Ext. Data Fig. 2b-c; Suppl. Table 1**). Large chromosomal rearrangements suggested by coverage analysis were all confirmed by pulse-field gel electrophoresis (PFGE) of yeast chromosomes, followed by long read Oxford Nanopore sequencing and *de novo* genome assemblies.

**Figure 2.**
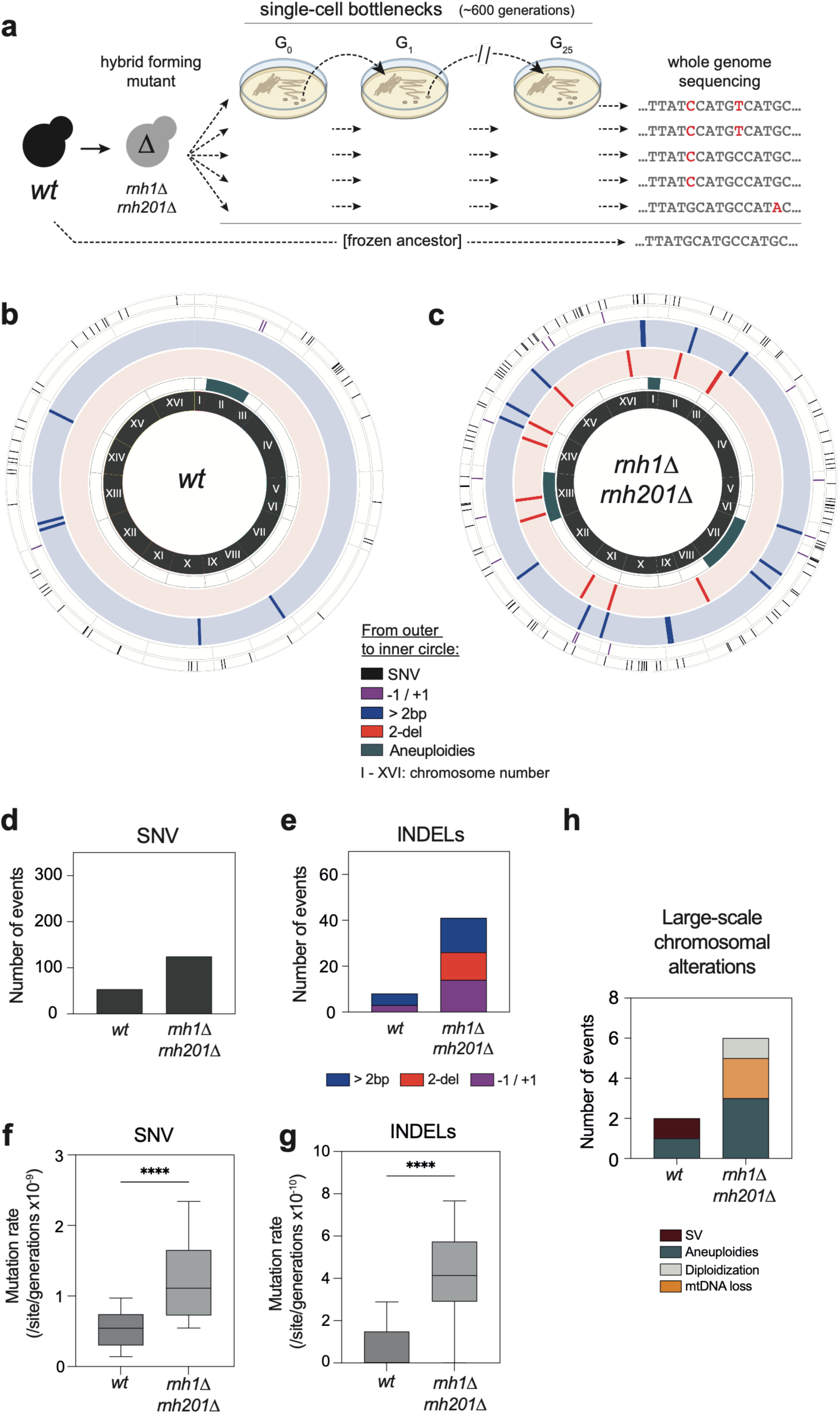
The mutational landscape of hybrid accumulation. **a**, Principle of the mutation accumulation line analysis. **b-c**, Circos plot representation of the genetic alterations identified in *wt* (**b)** and *rnh1Δ rnh201Δ* (**c**) MA lines. The positions of the indicated genetic alterations are shown relative to their chromosomal locations, with chromosomes arranged in a circular layout. **d-e**, Total number of SNVs (**d**) and INDELs (**e**) detected in MA lines either *wt* (n=16) or *rnh1Δ rnh201Δ* (n=15). The different types of INDELs are indicated. **f-g**, Mutation rates for SNVs (**f**) and INDELs (**g**) were calculated based on the number of genetic alterations detected per site per generation for each MA line (*wt*, n=16; *rnh1Δ rnh201Δ*, n=15). **h**, Total number of large-scale chromosomal alterations detected in *wt* and *rnh1Δ rnh201Δ* MA lines. The different types of alterations are represented. **** p<0.0001 (Mann-Whitney-Wilcoxon rank sum test).

Wild type cells mostly exhibited SNVs (**Fig. 2b**), with a base substitution rate estimated at 6 × 10^-10^ substitutions per site per generation (**Fig. 2f**), in agreement with previous reports^51–53^. Strikingly, RNase H-deficient cells exhibited a composite mutational signature that differed substantially from the wild type in both spectrum and burden (**Fig. 2c**). On the one hand, SNVs were markedly increased (two-fold) in mutant lines (**Fig 2d,f**), with no bias relative to *wt* regarding distribution along chromosomes or base substitution types **(Ext. Data Fig. 2d-g)**. On the other hand, INDELs exhibited a fourfold increase in the mutant **(Fig. 2e,g)**, with a mutational spectrum characterized by the emergence of a distinct class of 2 bp deletions (2-del), undetected in *wt* lines, together with an enrichment of other, longer INDEL events (**Fig. 2e**). Finally, other types of genetic alterations, mostly aneuploidies, were also increased in the mutant (**Fig. 2h**). In one line, spontaneous diploidization was observed (**Ext. Data Fig. 2b**), a common outcome in laboratory evolution experiments^54,55^ that is typically enhanced in mutator backgrounds^52^.

Strikingly, the increase in base substitutions and INDELs observed in hybrid-accumulating cells in lab settings (**Fig. 2d-g**) echoes the mutational signature scored in hybrid-prone genes within natural yeast populations (**Fig. 1g-h**). Overall, our data thus reveal a complex hybrid-associated mutational signature marked by an increase in base substitutions, the emergence of multiple classes of INDELs and the appearance of large-scale chromosomal alterations.

### Deconvolving the mutational signatures of ribonucleotide insertions vs. genic hybrids

The spectrum identified above likely results from mutagenesis induced by the combined accumulation of ribonucleotides and genic hybrids, both of them being increased upon simultaneous inactivation of the two RNases H (**Fig. 3a**, *top panel*). Supporting this idea, the 2 bp deletion signature was previously scored as the outcome of unfaithful ribonucleotide repair in the absence of RNase H2^20,56^. To further disentangle the respective contributions of both types of hybrids, we performed the same series of experiments in (i) a mutant of the THO complex (*hpr1Δ,* referred to as *tho*), a key RNA-binding complex preventing the accumulation of genic hybrids^33^ (**Fig. 3a**, *mid panel*) and (ii) the *rnh201-RED* (Ribonucleotide Excision Deficient; *P45D, Y219A*) separation-of-function mutant (**Fig. 3a**, *bottom panel*), which solely impacts RNase H2 ribonucleotide excision activity while retaining its hybrid removal function^57^. Both alleles were introduced in the same G_0_ *wt* background used for MA line generation, exhibited division rates comparable to *rnh1Δ rnh201Δ* cells (**Ext. Data Fig. 2a**), and displayed the expected specific hybrid accumulation phenotypes. Indeed, alkaline electrophoresis revealed extensive genomic DNA cleavage at embedded ribonucleotides in the *rnh201-RED* background, a phenotype undetected in the *tho* mutant (**Fig. 3b, Ext. Data Fig. 3a**). In contrast, dot blot analysis using the S9.6 antibody, which typically detects longer DNA:RNA hybrids, showed increased signals in the *tho* mutant, but not in the *rnh201-RED* background (**Fig. 3c-d**). Those hybrids were further mapped in *tho* cells by qDRIP, as above. Signals showed high correlation between biological replicates, sensitivity to *in vitro* RNase H treatment and strand-specificity over transcription units (**Ext. Data Fig. 3b-c; Fig. 3e**, *top panels*). Overall, THO inactivation detectably triggered hybrid accumulation at a limited number of loci (*e.g.* **Fig. 3e**, *top right panels*). We hypothesized that hybrids accumulating in these mutant cells would also be targeted by RNases H, as shown in *wt* cells (**Fig. 1**). To test this hypothesis, we mapped RNase H1 genome-wide binding sites in the same mutant background using H-CRAC (RNase H crosslinking and analysis of cDNAs)^16^. Both *wt* and *tho* mutant cells exhibited strand-specific binding of RNase H1 at the same genes where hybrids were detected by qDRIP in both backgrounds (**Fig. 3e**, *bottom panels*). Notably, RNase H1 binding was overall enhanced on a number of loci in the *tho* mutant (**Fig. 3e**, *bottom panels*). This supports the idea that the *tho* mutant background triggers the increased formation of transient hybrids which are further removed by RNases H, consistent with earlier observations^33,58^.

**Figure 3.**
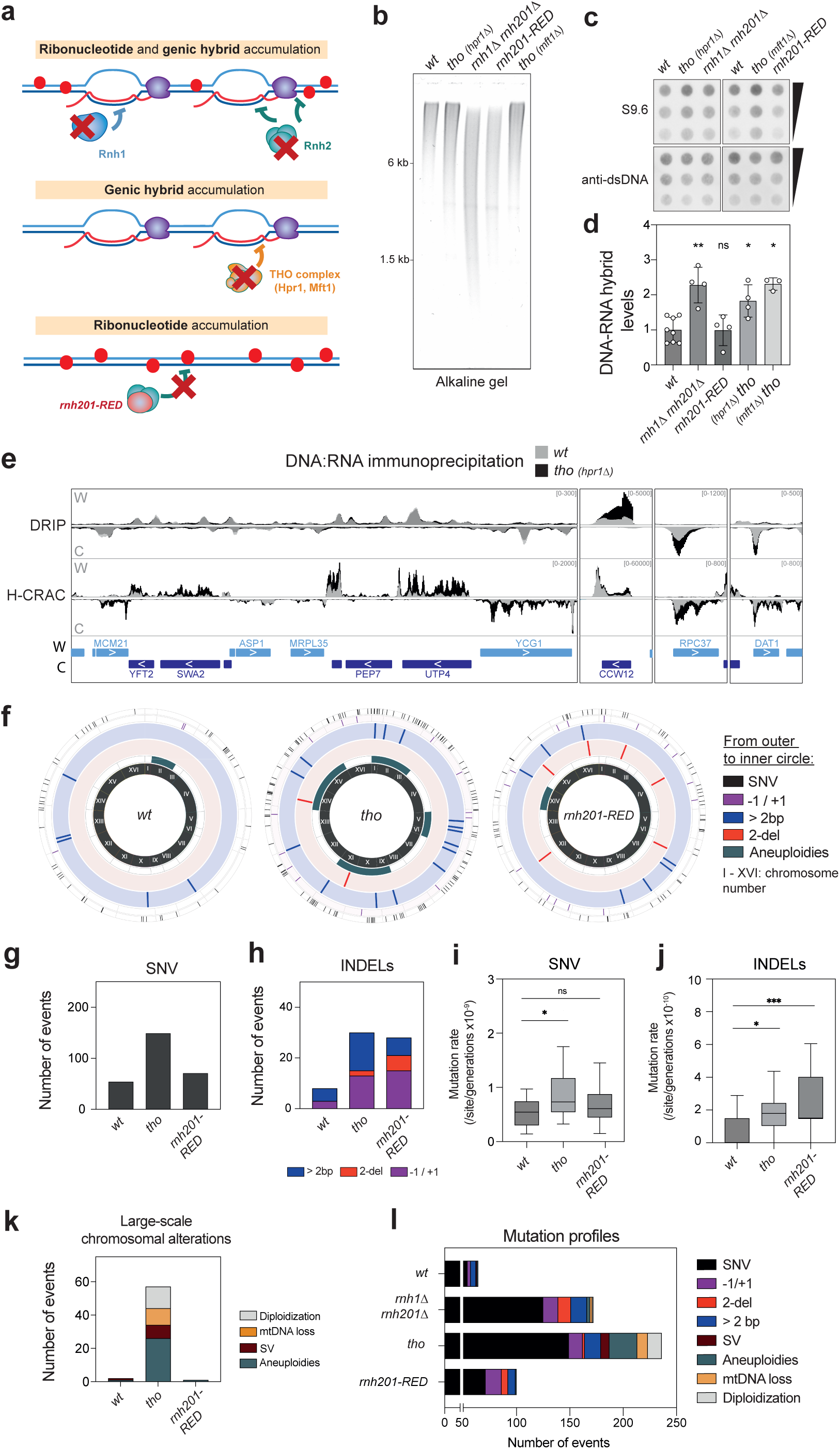
Distinct mutational spectra caused by ribonucleotides and genic hybrids. **a**, Principle of the separation of function approach. In the absence of RNase H1 and RNase H2 (*rnh1Δ rnh201Δ*; *top panel*), both genic DNA:RNA hybrids and embedded ribonucleotides (red dots) accumulate. In mutants of the THO complex (*mid panel*), only genic hybrids accumulate. In cells carrying the *rnh201-RED* allele (*bottom panel*), only ribonucleotides accumulate. **b**, Alkaline gel electrophoresis of genomic DNA from the indicated strains. Note the smear of cleaved DNA and the absence of high-molecular weight species in *rnh1Δ rnh201Δ* and *rnh201-RED* samples, indicating alkaline cleavage at embedded ribonucleotides. The position of molecular weights is indicated (kb). **c**, Dot blot analysis of DNA:RNA hybrid levels in genomic DNA samples from the indicated strains. DNA:RNA hybrids were detected with the S9.6 antibody (*top panel*) and dsDNA levels were used as loading control (anti-dsDNA, *bottom panel*). Decreasing amounts of genomic DNA were loaded to allow quantification. **d,** Quantification of dot blot analyses performed in **c** (mean±SD; n ≥ 3; relative to dsDNA and mean *wt* values). **e,** IGV representative screenshot of strand-specific coverage of DRIP-seq (*top panel*) and H-CRAC (*bottom panel*) in cells either *wt* (grey) or *tho (hpr1Δ,* black). Signals from Watson (W) or Crick (C) strands are represented. **f,** Circos plot representation of the genetic alterations identified in *wt* (same as Fig. 2b)*, tho (hpr1Δ)* and *rnh201-RED* MA lines. **g-h,** Total number of SNVs (**g**) and INDELs (**h**) detected in MA lines either *wt* (n=16), *tho* (*hpr1Δ*; n=16) or *rnh201-RED* (n=16). The different types of INDELs are indicated. **i-j**, Mutation rates for SNVs (**i**) and INDELs (**j**) in the indicated backgrounds (n=16 lines each). **k**, Total number of large-scale chromosomal alterations detected in the indicated MA lines. The different types of alterations are represented. **l**, Total number of genetic alterations in the different genetic backgrounds analyzed in this study. The different classes of genetic alterations are indicated. * p<0.05; ** p<0.01; *** p<0.001; ns, not significant (Mann-Whitney-Wilcoxon rank sum test).

Mutation accumulation experiments further revealed distinct patterns of genetic alterations in *tho* and *rnh201-RED* lines (**Fig. 3f)**. On the one hand, accumulation of genic hybrids (*tho*) was associated with an increase in both SNVs (**Fig. 3g,i**) and INDELs (**Fig. 3h,j**), as scored upon inactivation of RNases H, except for 2 bp deletions (**Fig. 2**). On the other hand, excess ribonucleotides were essentially associated with 2 bp deletion events (**Fig. 3h,j**), together with a slight increase in SNVs and -1/+1 deletions (**Fig. 3g-h**), this latter footprint being consistent with the role of embedded ribonucleotides in mismatch repair^59,60^. In both backgrounds, no bias in SNV subtype or genomic distribution were detected (**Ext. Data Fig. 2d-g**), as in *rnh1Δ rnh201Δ* cells. Finally, large-scale chromosomal alterations were specifically detected in *tho* cells, and consisted mainly of diploidization events and aneuploidies (**Fig. 3k**; **Ext. Data Fig. 2b-c**), as observed in *rnh1Δ rnh201Δ* cells, albeit to a higher extent. Of note, the enhanced frequency of diploidization did not confound the increased SNV rates, which were similarly elevated in haploid and diploid lines (**Ext. Data Fig. 3d**). However, diploidization likely facilitated the tolerance of structural variants, including heterozygous large-scale deletions and non-reciprocal translocations (**Fig. 3k, Ext. Data Fig. 3e**), that emerged in *tho* lines.

Overall, the mutational signature scored in the absence of RNases H (**Fig. 2**) can therefore be deconvolved into two components: (i) 2 bp deletions, uniquely recapitulated in *rnh201-RED* cells (**Fig. 3l**), a signature likely reflecting topoisomerase 1-mediated ribonucleotide excision in the absence of RNase H2^20,56^, and (ii) increased SNVs, INDELs and large-scale chromosomal alterations, which are shared by *tho* and *rnh1Δ rnh201Δ* cells and thereby attributable to genic hybrids. Defects in either genic hybrid prevention (by the THO complex) or removal (by RNases H) thus result in a common, composite and previously unrecognized mutational footprint on native genomes.

### DNA:RNA hybrids form genome-wide yet trigger instability primarily at repetitive DNA

To further evaluate whether the genetic alterations identified above arise from genic hybrids in *cis*, we compared the genomic distributions of mutations and hybrids, as measured in MA lines and qDRIP analyses, respectively. For this purpose, DRIP signals were represented as metaplots centered on mutation positions, both in *rnh1Δ rnh201Δ* and *tho* cells (**Fig. 4a-b**). In both cases, SNVs were not associated with DNA:RNA hybrid peaks (**Fig. 4a-b**, *black lines*), albeit enriched in transcribed regions in both mutants (**Ext. Data Fig. 4a**). This indicates that substitutions do not occur uniquely at the precise sites of hybrid accumulation, possibly resulting from longer ranges effect of hybrid processing and repair, *e.g.* through homologous recombination which involves DNA synthesis by error-prone DNA polymerases^61^ (see below, **Fig. 7c**). In contrast, INDELs (>2 bp) showed a marked colocalization with DNA:RNA hybrid peaks (**Fig. 4a-b**). This association was specific, as DRIP signals centered on randomly selected genomic positions (n=400) showed no enrichment in either *rnh1Δ rnh201Δ* or *tho* cells (**Fig. 4c**). These results thus support that hybrids accumulating upon THO or RNase H deficiency promote the formation of INDELs in *cis*.

**Figure 4.**
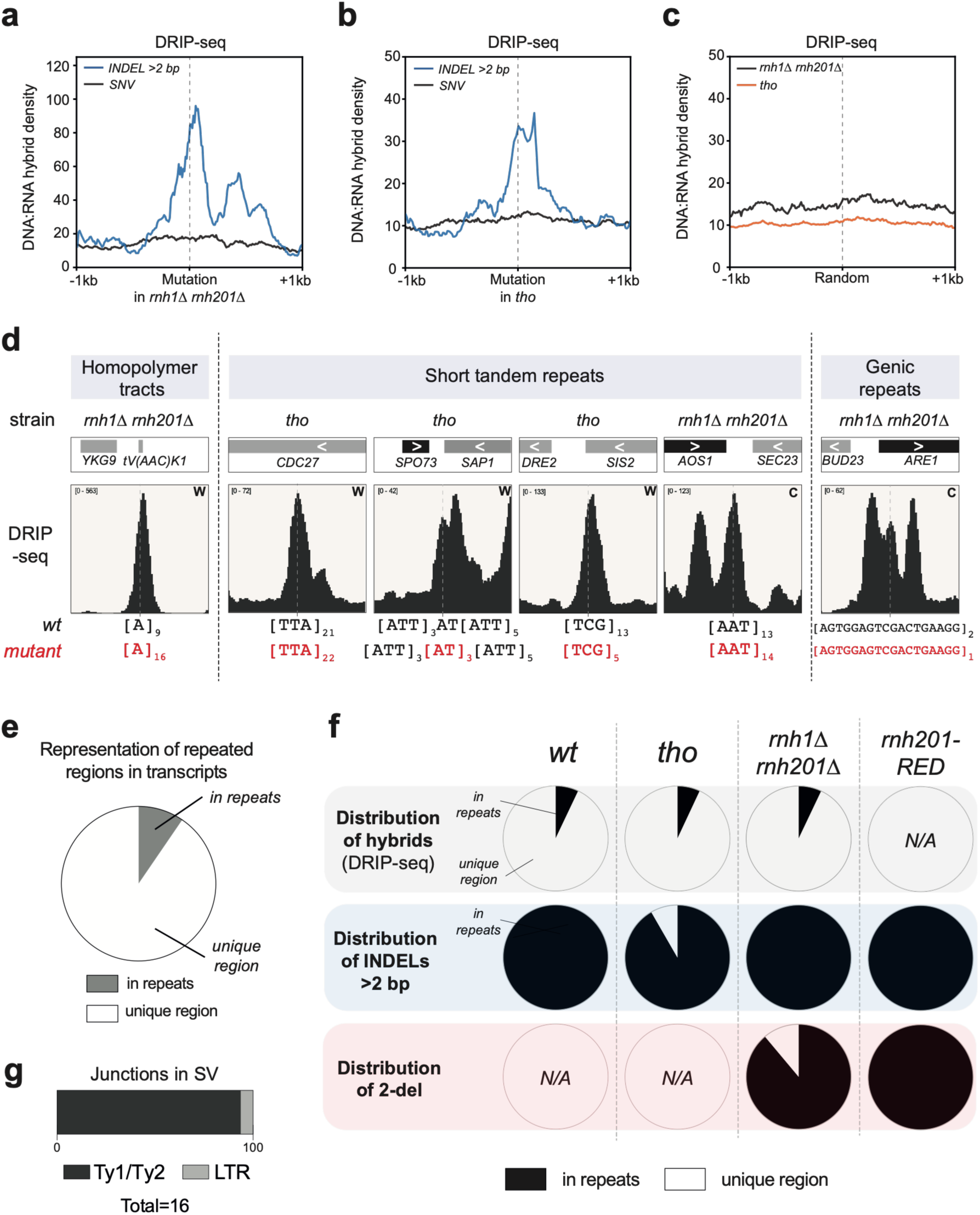
Hybrid genotoxicity is restricted to repetitive regions. **a**, Metaplot analysis of DNA:RNA hybrid densities in *rnh1Δ rnh201Δ* cells (DRIP-seq, from Fig. 1a) across regions centered on the indicated mutation types (SNVs, n=113, *black line*; INDELs, >2bp, n=14, *blue line*) detected in the same genetic background (Fig. 2c). **b,** Same as **a**, using DNA:RNA hybrid maps from *tho* (*hpr1Δ*) cells (Fig. 3e) and mutations (SNVs, n=129; INDELs, >2bp, n=13) identified in the same background (Fig. 3f, *mid panel*). **c**, Same as **a-b**, using DNA:RNA hybrid signals from *rnh1Δ rnh201Δ* (black line) or *tho* (orange line) cells, plotted across regions centered on 400 randomly selected genomic positions. Intergenic regions picked in the random sampling were excluded from the analysis. **d**, IGV representative screenshots showing DRIP-seq signals in the indicated strain (*rnh1Δ rnh201Δ* or *tho)* over genomic regions harboring typical INDELs identified in the corresponding MA lines. The sequences of the corresponding repetitive regions are shown for the parental G_0_ isolate (*wt*) and the derived (*rnh1Δ rnh201Δ* or *tho*) mutant lines. **e**, Representation of repetitive regions (as defined in **Methods**) within the transcribed subset of the genome. **f**, Distribution of the DNA:RNA hybrid signal (DRIP-seq; *top panel*) and INDELs (>2 bp and 2-del [2 bp deletions]; *middle and bottom panels*, respectively) across repetitive and unique genomic regions for the indicated genetic backgrounds. **g,** Localization of breakpoints in structural variants identified by long read sequencing in MA lines. LTR, long terminal repeat.

We noticed that DRIP peaks were slightly shifted downstream (3’) of mutation positions both in *rnh1Δ rnh201Δ* and *tho* cells (**Fig. 4a-b**), a pattern expected for INDELs occurring within repetitive sequences and being mapped at the 5′ boundary of the repeat tract. Consistently, most INDELs were found to occur in transcribed repetitive regions (**Fig. 4d**), including microsatellites, *i.e.* homopolymer tracts and short tandem repeats, as frequently present in 5’ or 3’ UTRs, or genic repeats encoding repeated protein domains (*e.g. ARE1*). At the genome level, only ∼10% of the transcribed regions correspond to curated repetitive sequences (**Fig. 4e**), including homopolymers (≥4bp) and tandem repeats. Hybrid-forming sites, as mapped across all our qDRIP datasets, also encompass a similar proportion of repetitive sequences (**Fig. 4f**, *top panel*). However, this specific fraction of the hybrid-forming genome concentrated most of the INDELs, including 2 bp deletions in *rnh201* mutants, or other type of INDELs in mutants accumulating genic hybrids (**Fig. 4f,** *mid and bottom panels*). We also analyzed the localization of large-scale genetic alterations, focusing on structural variants for which junctions were precisely determined by long read sequencing (**Ext. Data Fig. 4b-c**). Strikingly, all deletion and translocation breakpoints mapped to full-length Ty1 or Ty2 retroelements (6kb) or their long terminal repeats (LTR, 334 bp) (**Fig. 4g**), which are present in 32, 13, and 279 copies, respectively, in the yeast genome^62^, and represent a major source of non-allelic recombination^63^. Remarkably, the identification of repetitive elements as hotspots of genetic instability in hybrid-accumulating lab strains mirrors our observation of elevated mutagenesis at repetitive sequences within hybrid-prone genes in natural yeast populations (**Fig. 1h**). Interestingly, R-loop accumulating mutants were previously reported to exhibit genetic instability over triplet nucleotide repeat reporters^24,64^. Altogether, our data thus establish that while genic hybrids can accumulate genome-wide, their genotoxic effects are largely confined to repetitive regions in natural genomes.

### Mismatch repair and homologous recombination trigger genetic instability at natural sites of hybrid formation

Because mutational signatures from yeast populations and MA experiments are limited in size and difficult to investigate mechanistically, we developed dedicated assays to validate the observed genetic alterations and dissect their mechanisms of emergence. To assess SNVs and INDELs, we took advantage of the endogenous *LYS2* locus, a *bona fide* R-loop-forming gene^41,65^, which exhibits robust hybrid formation in both our DRIP and H-CRAC datasets (**Fig. 5a,** *top and middle panels*), contains repetitive sequences including short tandem repeats (**Fig. 5a**, *bottom panels*), and is amenable to forward mutation assays^66^. Using classical fluctuation analyses (**Fig 5b**, *left*), we observed increased mutation rates in both *rnh1Δ rnh201Δ* and *tho* strains (**Fig. 5c**), confirming their shared mutator phenotypes scored in MA lines analysis and linked to genic hybrid accumulation.

**Figure 5.**
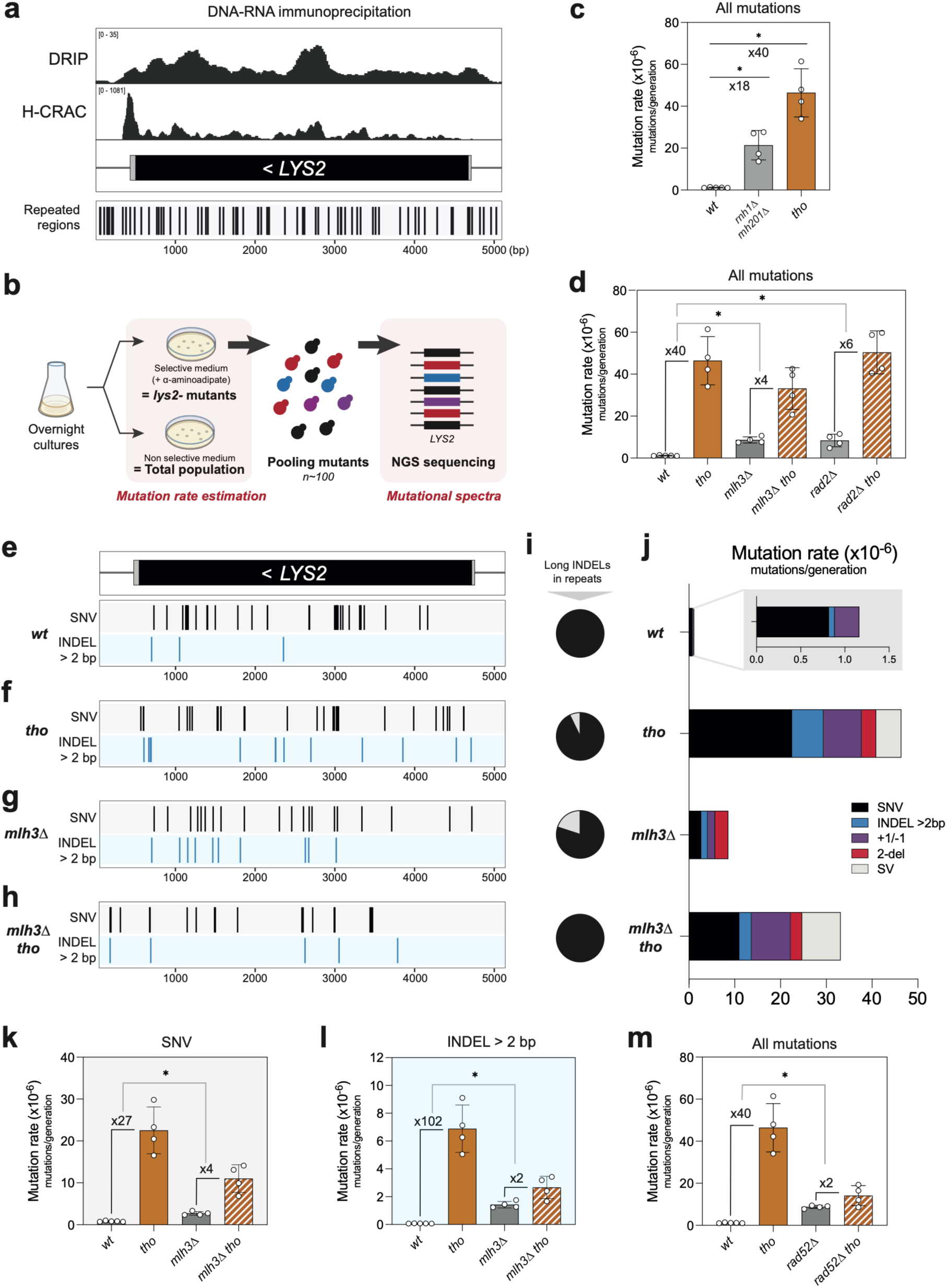
MutLγ and Rad52 drive hybrid-dependent SNVs and INDELs formation. **a**, IGV screenshot of DNA:RNA hybrids signals at the *LYS2* locus in *wt* cells, as measured by DRIP-seq (*top panel*) and H-CRAC (*mid panel*). The positions of repetitive regions within the *LYS2* genic sequence are indicated (*bottom panel*). **b**, Principle of mutation rate and mutation spectra determination for the *LYS2* locus. **c-d**, Mutation rates at the *LYS2* locus (mutations per generation, mean±SD, n=4) as measured in the indicated genetic backgrounds: *wt, tho (mft1Δ), rnh1Δ rnh201Δ, mlh3Δ, mlh3Δ tho (mft1Δ), rad2Δ, rad2 tho (mft1Δ)*. Note that the THO mutant *mft1Δ* was used instead of *hpr1Δ* in fluctuation assays because the latter exhibited reduced fitness on selective media (see **Methods**). **e-h,** Positions of SNVs and INDELs (>2bp) along the *LYS2* sequence in the same genetic backgrounds. This qualitative representation features the distribution of ∼100 mutations for each genotype, but does not account for the different mutation rates in each genetic background, as measured in panel **d**. **i**, Fraction of INDELs localizing to repetitive regions (in black) in the indicated genetic backgrounds. **j,** Mutation rates for the different types of genetic alterations identified in the indicated backgrounds. **k-l**, Mutation rates for SNVs (**k**) and INDELs (**l**) at the *LYS2* locus (mutations per generation, mean±SD, n=4) as measured in the same genetic backgrounds. **m**, Mutation rates at the *LYS2* locus (mutations per generation, mean±SD, n=4) as measured in the same genetic backgrounds. * p<0.05 (Mann-Whitney-Wilcoxon rank sum test). Note that the mutation rate values featured for *wt* and *tho (mft1Δ)* are identical across panels **c**, **d**, and **m**.

To define the mechanisms specifically underlying hybrid-dependent mutagenesis, we next focused on the *tho* mutant. To first assess the role of endonuclease-mediated cleavage, we performed the same assay upon inactivation of the nucleases previously proposed to target R-loop-forming regions based on reporter systems in yeast^22,24^. Inactivation of the Nucleotide Excision Repair (NER) nuclease Rad2^XPG^ or the Mismatch Repair (MMR) endonuclease MutLγ (*mlh3Δ*), a minor MutL complex with a limited role in replication-associated repair, both reduced *tho-*induced mutagenesis, with *MLH3* inactivation having the more pronounced effect (**Fig. 5d**). To assess the specific impact of this mechanism on the generation of SNVs and INDELs, we further analyzed mutational spectra by deep sequencing of the complete *LYS2* gene in pools of independent *lys2^-^* mutants (**Fig. 5b***, right*). Individual verification of *LYS2* amplicons followed by Sanger sequencing of a central region within the *LYS2* locus allowed to evaluate the fraction of *lys2*^-^ mutants corresponding to large-scale rearrangements, as previously scored in forward mutation assays^19^, and to validate deep sequencing analyses (**Ext. Data Fig. 5a-b**). Both SNVs and INDELs were detected, as exemplified by the profiling of ∼100 *lys2*^-^ mutants for each genotype (**Fig. 5e-h**). Of note, INDELs mostly occurred within repetitive regions (**Fig. 5i**), mirroring the mutation accumulation phenotypes observed in the complete yeast genome. Similarly, hybrid accumulation (*tho*) led to an increase in mutation rates for both SNVs (**Fig. 5j-k**) and INDELs (**Fig. 5j,l**), in line with MA lines observations. Strikingly, while single inactivation of Mlh3 (MutLγ) increased both types of mutations, in agreement with its reported supporting role in MMR during vegetative growth^67^, it markedly decreased both their frequencies in hybrid-accumulating conditions (compare *mlh3Δ* and *mlh3Δ tho*, **Fig. 5k-l**).

In view of the prominent role of endonuclease-mediated cleavage suggested by these experiments, we finally wondered whether hybrid-dependent breaks would be further repaired by homologous recombination, as previously shown in reporter assays^35^, possibly underlying the emergence of SNVs and INDELs. Loss of the central homologous recombination factor Rad52 itself promoted mutagenesis (**Fig. 5m**), in line with earlier observations that homologous recombination defects associate with increased mutation rates^46,68^. Remarkably, *RAD52* inactivation virtually suppressed the increased mutation frequencies caused by hybrid accumulation (**Fig. 5m**).

Altogether, our data thus indicate that homologous recombination is required for the emergence of both SNVs and INDELs, likely following Mlh3-dependent cleavage at hybrid-forming loci. Since MutLγ preferred substrates are DNA duplexes containing short extrahelical loops^69^, we propose that slipped-strand intermediates generated during the formation or resolution of hybrids on repetitive DNA constitute the primary substrates for MutLγ cleavage, thereby promoting genetic instability (see below, **Fig. 7c**).

### Long range effects of hybrids on genetic instability

To similarly assess the mechanisms driving structural variant formation upon hybrid accumulation, we designed a reporter strain to track a large-scale (40kb) deletion between two Ty1 elements on chromosome XVI, which was detected in one of our diploidized *tho* MA lines (**Ext. Data Fig. 4b**). For this purpose, we inserted in diploid strains the *URA3* and *HygroR* selection markers on each side of the downstream Ty1 element involved in the deletion (**Fig. 6a**). Deletion frequency was measured in fluctuation assays by counting the fraction of *ura3^-^ hyg^R^* mutants (**Fig. 6b**). In diploids, such mutants could carry either mutation(s) within the *URA3* gene or loss of the *URA3* locus by deletion of the entire chromosomal region located between the two Ty1 elements (**Ext. Data Fig. 6a***, top panel*). To estimate the contribution of these two mutational events in our assay, we analyzed in parallel the formation of *ura3^-^ hyg^R^* mutants in haploid lines in which *URA3* inactivation could solely arise from mutation at the *URA3* locus, since the 40kb chromosomal region lost in the deletion encompasses several essential genes (**Ext. Data Fig. 6a***, bottom panel*). Importantly, the rate of *URA3* inactivation was one order of magnitude lower in haploid that in diploid cells, supporting that our assay scores deletion events in diploids, as further confirmed by PFGE analysis (**Ext. Data Fig. 6b-c**).

**Figure 6.**
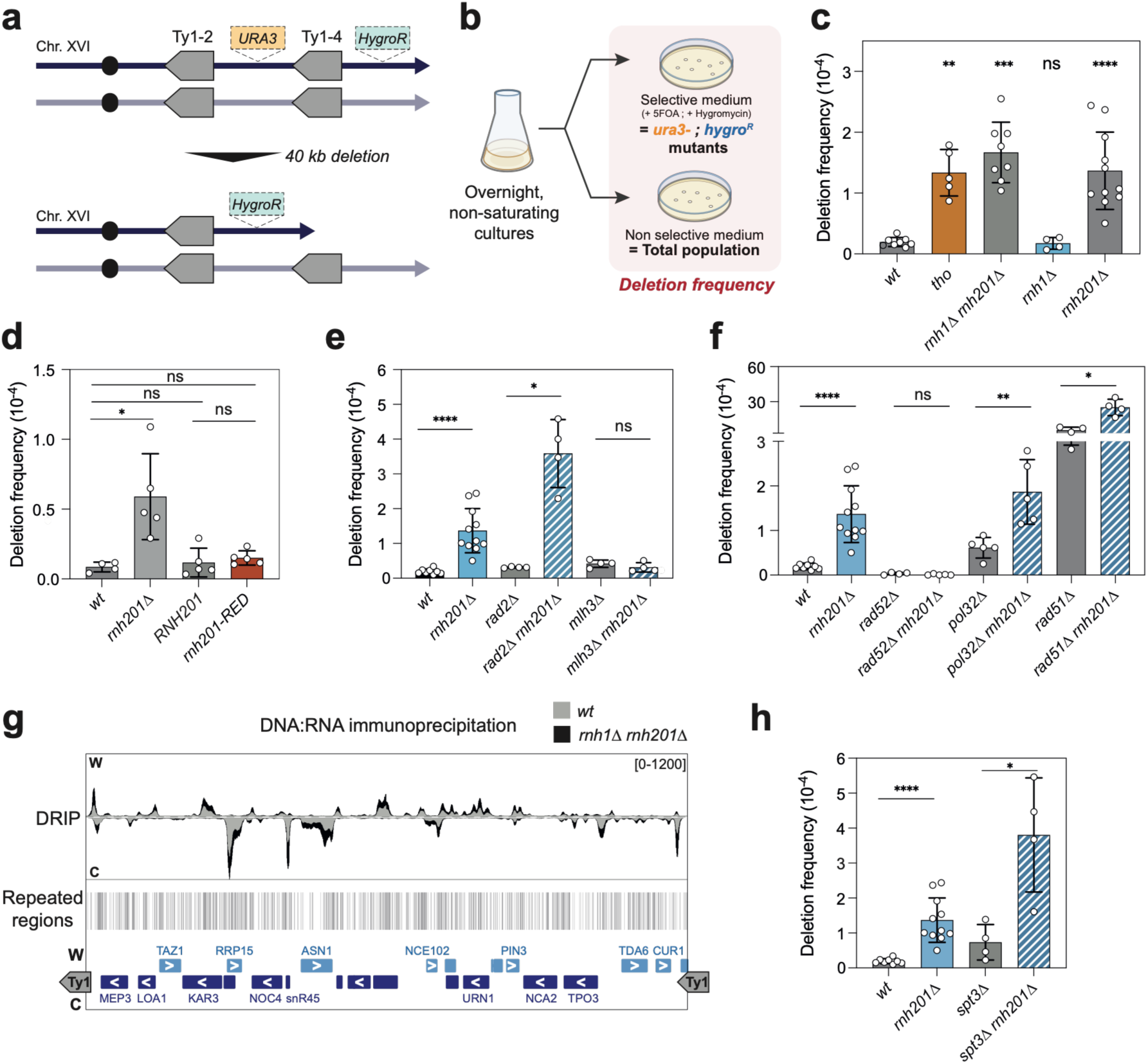
MutLγ and Rad52 drive hybrid-dependent genome rearrangements. **a**, Principle of the chromosomal reporter system used to score deletions in between Ty1-2 and Ty1-4 retroelements on chromosome XVI. **b,** Principle of the deletion reporter assay. **c-f**, Deletion frequencies (*ura3^-^ hyg^R^* fraction, mean±SD, n ≥ 4), as measured in the indicated genetic backgrounds. *tho* refers to the *mft1Δ* mutant. *RNH201* and *RNH201-RED* backgrounds were obtained by transforming *RNH201* and *rnh201-RED* plasmids in *rnh201Δ* cells carrying the deletion reporter. **g**, IGV screenshot of DNA:RNA hybrids signals within the 40kb chromosome XVI region located in between Ty1-2 and Ty1-4, as measured by DRIP-seq in *wt* and *rnh1Δ rnh201Δ* cells (*top panel*). The positions of repetitive regions are indicated (*bottom panel*). **h**, Deletion frequencies, as in **c-f.** * p<0.05; ** p<0.01; *** p<0.001; **** p<0.0001; ns, not significant (Mann-Whitney-Wilcoxon rank sum test). Unless otherwise indicated, all statistical comparisons were made against *wt*.

With this system, we first confirmed increased deletion rates in *tho* cells (**Fig. 6c**), a phenotype also observed upon RNase H inactivation (*rnh1Δ rnh201Δ*) and largely attributable to loss of *RNH201* (**Fig. 6c**). To determine whether this reflected excess ribonucleotide incorporation rather than DNA:RNA hybrid accumulation, we complemented *rnh201Δ* cells with either wild-type or ribonucleotide excision-deficient (*RED*) *RNH201* alleles. Both fully suppressed the deletion phenotype (**Fig. 6d; Ext. Data Fig. 6d**), demonstrating that DNA:RNA hybrids, rather than ribonucleotides, are the common trigger of these large-scale deletions.

To investigate the mechanisms underlying hybrid-dependent deletion, we inactivated the R-loop-associated endonucleases Rad2^XPG^ and MutLγ (*mlh3Δ*), as above. Strikingly, while *RAD2* loss had little effect, inactivation of MutLγ (*mlh3Δ*) completely suppressed the deletion caused by hybrid accumulation (**Fig. 6e**). To determine whether Mlh3-dependent cleavage at hybrid-forming loci promotes recombination, as shown above (**Fig. 5**), we disrupted distinct homologous recombination pathways in our deletion system. Whereas *RAD52* inactivation completely suppressed hybrid-induced deletions, absence of either Rad51, required for homology-directed repair, or the BIR factor Pol32 had no effect on the stimulatory effect of *RNH201* inactivation (**Fig. 6f**). Together, these findings indicate that MutLγ-dependent cleavage promotes Rad52-dependent genome rearrangements, likely through SSA between homologous Ty1 elements.

Mlh3-dependent cleavage could initiate rearrangements either within the ∼40-kb interval separating the two Ty1 elements on chromosome XVI or within the Ty1 elements themselves. qDRIP identified multiple RNase H-sensitive hybrid-forming loci across the intervening region (**Fig. 6g**), and DRIP analysis following restriction digestion further detected genomic DNA:RNA hybrids at both Ty1 elements (**Ext. Data Fig. 6e**), while excluding reverse-transcription intermediates^70^. To discriminate the source of genotoxic hybrids in this context, we specifically interfered with the formation of Ty1 genomic hybrids by inactivating *SPT3*, a SAGA subunit strictly required for Ty1 transcription^71^. Although *SPT3* deletion strongly reduced Ty1 expression (**Ext. Data Fig. 6f**), it did not suppress the increased deletion rate of *rnh201Δ* cells (**Fig. 6h**), indicating that hybrids forming between Ty1 elements, rather than within them, are the primary source of genome instability. Consistent with this conclusion, the intervening region contains multiple hybrid-forming repetitive sequences (**Fig. 6g**), which likely represent the initial sites of MutLγ-dependent cleavage. Genic hybrids can therefore compromise genome stability beyond their site of formation, promoting genetic alterations at distant loci.

### A model for hybrid-dependent genotoxicity in a native genome

Based on our reporter analyses, the three classes of genetic alterations that define the mutational signature of hybrid genotoxicity in MA lines – *i.e.*, SNVs (**Fig. 5k**), INDELs (**Fig. 5l**), and large-scale chromosomal rearrangements (**Fig. 6e**) - share a common genetic dependency on MutLγ, which likely acts primarily at repetitive DNA, and Rad52. This finding challenges the prevailing view that hybrid-associated genome instability arises predominantly from transcription–replication conflicts (TRCs), a model largely supported by reporter-based studies in bacteria, yeast, and mammalian cells, demonstrating that head-on (HO) collisions between transcription and replication fork are more deleterious than their codirectional (CD) counterparts^26,27,72^. To assess whether TRCs contribute to the mutational signature of hybrids in the native yeast genome, we analyzed the positions of genic SNVs and INDELs scored in MA lines with respect to the orientation of replication, based on published OK-seq data^73^. Genes transcribed in HO and CD orientations relative to replication were equally represented in the whole yeast genome (**Fig. 7a**). Remarkably, both types of genetic alterations (SNVs and INDELs) were similarly represented in both TRC orientations, in *wt* and hybrid-accumulating conditions (compare HO vs. CD, **Fig. 7b**). Altogether, our data thus suggest that in the natural yeast genome, TRCs are not the main contributors to genome instability upon hybrid accumulation, as recently proposed based on mutation accumulation in bacteria^37^. Instead, our data support a mechanism in which MutLγ-dependent cleavage is the primary initiating event in the mutagenic processing of hybrid-associated lesions at repetitive DNA, stimulating Rad52-dependent recombination events (**Fig. 7c**).

**Figure 7.**
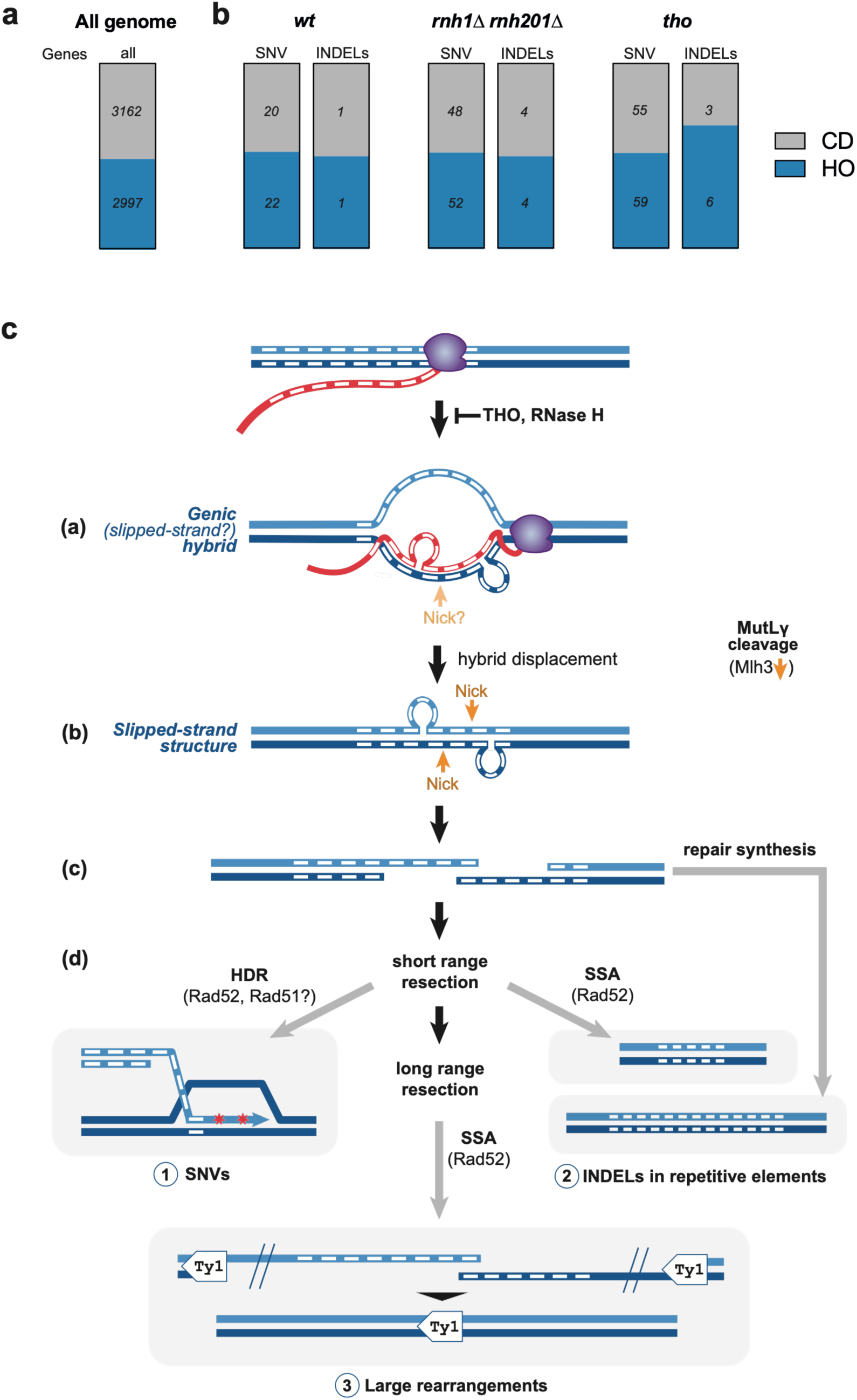
A model for hybrid-induced genetic instability in a native genome. **a,** Proportions of protein-coding genes transcribed in a co-directional (CD) or head-on (HO) orientation relative to replication. **b**, Proportion of SNVs and INDELs occurring in genes transcribed in CD or HO orientation relative to replication, in the indicated MA lines. The number of each type of genetic alteration is indicated for the different categories. **c**, Model for hybrid-dependent genetic instability arising at repetitive sequences, such as homopolymeric tracts or trinucleotide repeats (represented by white rectangles). During R-loop formation (a), or following hybrid displacement by helicases or RNases H (b), slipped-strand structures may form at repetitive sequences, generating extruded loops within the DNA:RNA hybrid (a) or the re-annealed DNA duplex (b). These slipped-strand structures may be further stabilized by secondary structures, *e.g.* hairpins (not depicted), formed by repeats within the extruded DNA or RNA loops. MutLγ-dependent cleavage at these structures (orange arrows) would then generate single-strand breaks (c). Following short-range resection, homology-directed repair may restore the damaged region, although nucleotide substitutions (SNVs) could be introduced during repair synthesis (d, 1). Alternatively, single-strand annealing (SSA) may generate deletions within repetitive sequences (d, 2). Following extensive resection, homeologous sequences, such as Ty1 elements, may become exposed, promoting SSA-mediated large-scale genome rearrangements, including deletions (d, 3). Repeat expansions may also arise through repair synthesis of the double-nicked intermediate, as previously proposed for MutLγ-dependent cleavage events^69^.

## Discussion

By analyzing in parallel genetic variation in hybrid-prone genes across natural yeast genomes and *de novo* mutations accumulating under laboratory conditions, we revealed a composite mutational footprint comprising SNVs, INDELs and large-scale chromosomal rearrangements, that recurrently emerges in distinct contexts of hybrid formation. While the three classes of genetic alterations defining this footprint can also arise in other mutator contexts^46,52,74^, their recurrent emergence as a consequence of hybrid accumulation constitutes the defining feature of the hybrid-associated mutational signature. SNPs and point mutations are enriched at hybrid-forming genes in *wt* populations and at the hybrid-prone *LYS2* locus, yet they do not precisely coincide with DNA:RNA hybrid maxima, suggesting that they arise during the recombinogenic repair of hybrid-associated lesions and therefore occur at a distance from the initiating hybrid. In contrast, INDELs precisely map at DNA:RNA hybrid peaks within repetitive sequences, consistent with hybrid-forming repeats acting as the primary lesion. Large-scale chromosomal alterations likewise exhibit breakpoints proximal to DNA:RNA hybrids, as exemplified by the chromosome XVI deletion.

Although this signature strikingly differs from that reported upon RNases H inactivation in bacteria, which mostly revealed footprints of inserted ribonucleotides but modest effects of genic DNA:RNA hybrids^37,75^, it is consistent with the spectrum of genetic alterations previously scored in yeast RNase H2 mutants^56,76,77^, although absence of hybrid mapping precluded assessment of their direct roles in mutagenesis in these studies. The convergence of mutational outcomes following disruption of distinct hybrid regulators - THO/TREX; RNases H - further suggests that common downstream processing pathways act on similarly localized, transient hybrid structures. Indeed, hybrids accumulating upon THO/TREX inactivation can be efficiently removed by RNase H, in line with earlier studies^33,58^, paralleling the proposed role of the Sen1 helicase in resolving hybrids that arise in RNase H-deficient cells^29^. Nevertheless, distinct hybrid regulators may also suppress specific mutational footprints, as suggested here by the prominent role of RNase H2 in preventing Ty1-associated deletions or the more marked role of the THO/TREX complex at hybrids in specific genomic regions.

A striking conclusion of our study is that, despite their widespread formation, the genotoxic effects of DNA:RNA hybrids are largely confined to repetitive DNA. Repetitive sequences comprise a substantial fraction of eukaryotic genomes and are well-established hotspots of genome instability because of their susceptibility to replication errors and error-prone repair^78,79,52,80^. Of relevance here, artificial reporters or model genes containing transcribed repetitive elements were previously reported to exhibit hybrid formation and hybrid-dependent genetic instability^24,81–84^. Based on our genome-wide observations, we do not favor a mechanism in which repetitive DNA is intrinsically more prone to hybrid formation, as repeats are similarly represented in the genome and its hybrid-forming fraction. Instead, our data support that repetitive DNA is particularly susceptible to the mutagenic processing of hybrids, as previously proposed^85^.

Mechanismwise, hybrid formation or removal could promote the generation of slipped-strand structures at repetitive loci, creating substrates for MutLγ-dependent cleavage and Rad52-dependent recombination, whose activities appear central to the three types of mutational outcomes described here. Consistently, MutLγ preferably engages on slipped-DNA intermediates and has previously been implicated in repeat instability^69,86^. In the context of hybrid-forming loci, MutLγ could target slipped-strand DNA duplexes formed upon RNA removal by RNases H or helicases (**Fig. 7c**, (b)), consistent with earlier observations^24,84,87^. Alternatively, MutLγ could directly recognize slipped-strand DNA:RNA hybrid duplexes formed at hybrid-forming repeats (**Fig. 7c**, (a)), although this hypothesis awaits biochemical investigation. Depending on subsequent repair, the resulting breaks could generate local mutations within repetitive DNA or more distant genome alterations through allelic or non-allelic recombination, including the Ty1-mediated deletions described here (**Fig. 7c**).

While Rad52 is essential for the different types of mutations scored here, loss of MutLγ does not fully suppress SNVs and INDELs generation, suggesting the existence of other nucleases at play at hybrid regions. Of note, hybrid-associated genetic alterations are not detectably enriched at head-on genes (**Fig. 7b**), suggesting that they do not arise from conflicts with replication, in agreement with previous studies using repeat-containing reporters^83,84^. Overall, our proposed model echoes the mechanism underlying ribonucleotide-associated mutagenesis, as also scored here, in which cleavage at embedded ribonucleotides by topoisomerase 1 is followed by error-prone repair in dinucleotide repeat contexts^20,56^. Together, our findings thus identify repetitive DNA as a major substrate through which distinct classes of DNA:RNA hybrids lead to genetic instability.

Identifying mutational signatures associated with specific DNA metabolic processes is essential for understanding their mechanisms and contribution to human disease. As illustrated by ribonucleotide-associated mutagenesis, signatures first identified in model systems can subsequently be recognized in human cells and tumors^88^. Likewise, DNA:RNA hybrids have been linked to cancer-associated INDELs and SVs^89–91^, yet hybrids have rarely been mapped in the same biological contexts, and many cancer mutation datasets are derived from exome sequencing^92,93^, biasing mutation detection toward transcribed, hybrid-prone regions. Genome-wide analyses integrating DNA:RNA hybrid maps with mutational landscapes will therefore be critical for establishing the contribution of hybrids to genome plasticity in physiological and pathological contexts, shedding light on their dual role in influencing genetic stability and driving genome evolution.

## Methods

### Yeast Strains and Growth

All *S. cerevisiae* yeast strains used in this study (listed in **Suppl. Table 2**) are isogenic to S288C, unless specified, and were obtained by homologous recombination and/or successive crosses according to standard procedures. Cells were grown at 30°C in standard yeast extract peptone dextrose (YPD) or synthetic complete (SC) medium supplemented with the required nutrients. For selection of *ura3* and *lys2* mutants, synthetic media included 5-fluoroorotic acid (Euromedex, 1g/L) and alpha-aminoadipate (Bachem, 2g/L), respectively^94^. Since *hpr1Δ* THO complex mutants exhibited decreased fitness in these two selective media, we used instead another mutant of the same complex (*mft1Δ*) with similar phenotypes yet milder growth defects^41,95^ for the corresponding fluctuation assays. Plasmids (listed in **Suppl. Table 3**) were used as templates for genome editing or for transformation.

### DNA:RNA hybrid detection and bioinformatic analysis

DNA:RNA hybrid immunoprecipitation coupled to sequencing (qDRIP) was performed according to a published procedure^47^, with our previously described modifications^41^. Briefly, 100 OD of cells were mixed prior to lysis with a synthetic DNA:RNA hybrid spike-in (1:1 ratio; **Suppl. Table 4**) and *hpr1Δ C. glabrata* cells (95:5 ratio; **Suppl. Table 2**). Genomic DNA was isolated by phenol-extraction and ethanol precipitation, fragmented in a Covaris focused-ultrasonicator (Covaris M220), treated or not with RNase H (New England Biolabs), and mixed 1 h at 4°C with 5.25 µg of S9.6 purified antibody in 20 mM Tris, 150 mM NaCl, 0.1% Tween-20, pH 8 (Tris-Buffered Saline [TBS]-Tween). Immunoprecipitation was achieved using Dynabeads Protein G (Thermo Fisher Scientific). Input and immunoprecipitated DNA were purified with the QIAquick DNA purification kit (Qiagen) and libraries were created using the xGen ssDNA & Low-Input DNA Library Prep Kit (Integrated DNA Technologies), with 16 cycles of amplification. Paired-end sequencing was achieved on an Illumina NovaSeq 6000 platform (Novogene). Raw paired-end reads were trimmed with Trimmomatic v0.39 to remove Illumina adapters (first 20 bases of each read), and low-quality bases (leading/trailing quality threshold = 3; sliding window of 4 bp with a minimum average Phred score of 15). Reads shorter than 36 bp were discarded, and only paired reads passing quality filtering were retained for downstream analyses, and further aligned to the SacCer3 (*S. cerevisiae*) and CBS138 (*C. glabrata*) genomes using Bowtie2 (version 2.5.1). Aligned reads were filtered to retain only the reads mapped in a proper pair with a quality score ≥30 using Samtools (version 1.8), and duplicates were removed using Picard (version 2.23.5). For *S. cerevisiae*, aligned reads were split based on their strand of origin using BamCoverage (version 3.5.4), with a bin size of 20 bp, no scaling, and only including reads originating from fragments from the Watson or Crick strands. Calibration was achieved using an occupancy ratio defined for each sample based on the number of *C. glabrata* reads in input and immunoprecipitated samples, as previously described^96^. Metaplots and heatmaps were generated using deepTools^97^ using previously described transcript annotations^98^.

DRIP-qPCR was performed as previously described^41^, except that genomic DNA was fragmented with a modified cocktail of restriction enzymes (BpiI, Eam1105I, Eco72I, Hin1I, NcoI and SmiI; FastDigest, Thermo Fisher Scientific), in order to generate restriction fragments encompassing YPRCTy1-2 and YPRCTy1-4 loci and their flanking genomic regions in chromosomes XVI. Input and immunoprecipitated DNAs were quantified by real-time PCR with a LightCycler 480 system (Roche) using SYBR Green incorporation according to the manufacturer’s instructions. % of immunoprecipitation (IP) were normalized to the % of IP obtained for the synthetic spike-in, resulting in adjusted % of IP.

Dot blot detection of DNA:RNA hybrids was performed as previously described^41^. Decreasing amounts of genomic DNA extracted as above were spotted on nylon membranes (Hybond-N+, GE Healthcare). Membranes were incubated with anti DNA:RNA hybrids (S9.6, Kerafast; 0.3 µg/mL in TBS, 0.5% Tween-20, 5% skimmed milk) or anti double-stranded DNA (HYB331-01, Santa Cruz Biotechnology; 1:5,000 in TBS, 0.5% Tween-20, 5% BSA) and revealed using chemiluminescent reagents (Supersignal, Thermo Fisher Scientific) following incubation with anti-mouse peroxidase-conjugated antibodies (Jackson ImmunoResearch). Relative DNA:RNA hybrid and total DNA amounts were quantified with ImageJ using images acquired through a ChemiDoc MP Imaging System (Bio-Rad), with serial dilutions of a reference sample as a standard.

RNase H cross-linking and analysis of cDNAs (H-CRAC) was performed as reported^16^. Briefly, cells expressing a His-TEV-proteinA-tagged version of RNase H1 from its genomic locus were crosslinked by UV exposure, cryogenically ground and RNA protein complexes were isolated by tandem affinity purification, with adaptor ligation carried on the final Ni-NTA column. Isolated RNAs were then reverse transcribed, PCR amplified and the cDNAs were sequenced on a NextSeq 500 Illumina sequencer. H-CRAC data analysis was performed as described, with normalization based on *S. pombe* spike-in cells^16^.

### Alkaline electrophoresis detection of ribonucleotides

Alkaline hydrolysis and alkaline agarose electrophoresis of genomic DNA was performed according to a published procedure^99^. Briefly, yeast spheroplasts were prepared by treatment with 0.5 mg/mL zymolyase 100T (MP Biomedicals) in 0.1M KPi pH7.5, 1.2M sorbitol, 0.1M EDTA, resuspended in 10 mM Tris pH 8, 0.1 M NaCl, 2% Triton X-100, 1% SDS, 1 mM EDTA, and genomic DNA was isolated using phenol-chloroform extraction and ethanol precipitation. DNA preps were treated with 0.7 mg/mL RNase A (Sigma) and 150 U/mL RNAse T1 (Applied Biosystems) and reprecipitated with ethanol. Either KOH or KCl (0.1M final) was added to 10 µg of genomic DNA for 2 h at 55°C. KOH-treated and KCl-treated samples were supplemented with 6X Alkaline and 6X Neutral loading buffer, respectively, and electrophoresis was achieved in either alkaline or neutral buffers, as described^99^. Gels were stained with SYBR Gold (Thermo Fisher Scientific; 1:10,000).

### Transcriptome analysis

For RNAseq, total RNAs were purified from 10 OD of cells using the Nucleospin RNAII kit (Macherey Nagel) according to the manufacturer’s instructions. Directional libraries were prepared with rRNA depletion and paired-end sequencing was achieved on an Illumina NovaSeq X Plus platform (Novogene). Differential expression analysis was achieved using DESeq2, as previously reported^41^.

For RT-qPCR, RNAs were similarly extracted and reverse-transcribed using random hexamers (P(dN)6, Roche) and Superscript II reverse transcriptase (Thermo Fisher Scientific). cDNAs were quantified by real-time PCR as above, using primers described in **Suppl. Table 4.**

Population-scale RNA abundance data were obtained from the RNA-seq dataset of 969 Saccharomyces cerevisiae isolates available through the 1002 Yeast Genomes Project (file final_data_annotated_merged_04052022.tab at http://1002genomes.ustrasbg.fr/files/RNAseq/), which provides normalized expression levels as transcripts per million (TPM)^50^. TPM values were extracted for hybrid-prone genes (n=71) and hybrid-poor genes (n=80) using their systematic gene names. For each gene, the mean, median, standard deviation (SD), coefficient of variation (CV = SD/mean), and the number of isolates with available TPM values were calculated. A gene-by-strain expression matrix containing TPM values for all 969 isolates was generated, and data processing and summary statistics were performed in Python using the pandas library.

### Variant analyses in yeast populations

Genetic variation was analyzed using a variant dataset comprising 3,570 Saccharomyces cerevisiae isolates from 40 published studies (doi.org/10.5281/zenodo.21238234). Variants were called and filtered from short-read sequencing data following GATK Best Practices recommendations. Repetitive regions, representing approximately 8% of the reference genome, were excluded. The resulting dataset contained 3.61 million SNPs and small insertions and deletions (INDELs). Analyses focused on 71 hybrid-prone genes and 80 hybrid-poor genes. For each group of genes, SNPs and INDELs overlapping these genes were extracted separately from the genome-wide VCF using bcftools and Python scripts based on gene coordinates. Variant effects were annotated using SnpEff v5.2, allowing SNPs to be classified as synonymous or nonsynonymous and INDELs as in-frame or frameshift^100^. For genome-wide analyses, we used bedtools intersect to compute variant counts in 100-bp genomic and to extract the 20 bp flanking sequences around INDELs. For PCA analysis, SNPs were filtered from the VCF file for biallelic SNP with a minor allele frequency > 5% and pruned for linkage disequilibrium (window = 50 bp, step = 1, r² ≤ 0.5).

### Mutation accumulation line generation and analysis

Mutation accumulation lines were generated according to a published procedure^52^. All relevant alleles (*rnh1Δ rnh201Δ, hpr1Δ, rnh201-RED*) were freshly introduced into a single-colony isolate of the parental *wt* strain (BY4741). At each bottleneck, cells from a single colony were streaked onto YPD agar plates and incubated for 3 days at 30°C. A randomly selected, average-sized unique colony was then used to initiate the next passage. For each genotype, 15-16 independent MA lines were propagated through 25 serial bottlenecks. Ploidy was checked by flow cytometry following propidium iodide staining of G_0_, G_25_ and intermediate isolates. The number of generations per bottleneck was estimated for each genotype by determining the number of cells per colony.

For short read sequencing, total genomic DNA was extracted from spheroplasts and treated with RNase A/T1 as described above, prior to whole genome sequencing. Paired-end libraries were prepared and sequenced at high coverage (>250X) on a DNBSEQ-G400 (BGI) or NovaSeq X Plus (Novogene) platforms. Paired-end reads were aligned to the S288C reference genome (R64-1-1_20110203) using BWA-MEM (v0.7.17). Duplicates reads were marked with Picard MarkDuplicates (v2.18.29), and variants were called using GATK HaplotypeCaller (v4.0.5.1). Raw variant calls were filtered using stringent hard-filtering criteria (QUAL ≥ 30, QD ≥ 6, MQ ≥ 59, FS ≤ 5, SOR ≤ 1.5, MQRankSum ≥ −1.5 and ReadPosRankSum ≥ −4), and clusters of variants occurring within 10 bp were removed. Mitochondrial variants and variants with insufficient read depth (the maximum between 5 or 25% of the mean sample depth) were excluded. To retain only lineage-specific mutations, variants detected in ancestral (G_0_) populations or shared between independent evolved lines at the final stage (G_25_) were removed using BEDTools (v2.31.0). VCF processing, filtering and summary statistics were performed using BCFtools (v1.17), SAMtools (v1.17) and VCFtools (v0.1.16).

For long read sequencing, genomic DNA was extracted either with QIAGEN genomic TIP 100/G or as previously described^101^. Size selection was performed using Circulomics SRE kit (PacBio). DNA library was prepared using either ONT LSK109 coupled with EXP-NBD104 or SQK-NBD114.24 kit from Oxford Nanopore. Sequencing was performed either on a MK&C Minion sequencer with R9.1 flowcell or on a P2solo sequencer using R10.4.1 flowcell. For the samples sequenced with R9 technology, basecalling was performed using Guppy (V6.1.5,model=dna_r9.4.1_450bps_hac.cfg) and barcode were removed with Porechop. For the samples sequenced with R10 technology, basecalling and barcode removing was performed with Dorado (V1.1.O, model=dna_r10.4.1_e8.2_400bps_sup@5.2.0). Sequences were downsampled using Filtlong to obtain a coverage of 40X. Genome assembly was performed using 3 different assemblers: Canu (v2.2)^102^, SMARTde-novo^103^ and NextDenovo^104^. Contigs were polished using 1 to 3 rounds of Racon (v1.4.21)^105^ and 2 rounds of Medaka (https://github.com/nanoporetech/medaka v1.2.3). Scaffolding was performed with Ragout (v2.3)^106^.

Mutational spectra were defined from short-read sequencing data by counting SNVs and short INDELS (<100bp). INDELs were further classified according to their size: -1/+1bp; 2bp deletions (2-del); >2bp (including 2 bp insertions). Aneuploidies were identified by genome-wide coverage analysis. Structural variants initially inferred from coverage analysis were mapped by aligning long read sequences on reference genome using Tablet (https://github.com/cropgeeks/tablet) to identify reads validating SVs junctions. Diploidization events were scored based on flow cytometry and mitochondrial DNA loss was evaluated by both coverage analysis and DAPI staining. All genetic alterations were represented along the yeast genome with CIRCOS plots^107^. Associations with repetitive sequences were assessed by intersecting variant coordinates with a curated map of *S. cerevisiae* repetitive elements including homopolymer stretches (≥ 4 nt) and low complexity repeats, simple stretches and simple repeats from the R64 annotation (https://hgdownload.soe.ucsc.edu/hubs/GCF/000/146/045/GCF_000146045.2/bbi/).

### Pulse-Field Gel Electrophoresis

Yeast DNA was extracted in agarose plugs, prepared according to a published procedure^108^ and sealed in a 1% Seakem GTC gel (1% aragose, 0.5x TBE). Migration was performed in a CHEF-DRIII (BioRad) system with the 2 steps program (60s switching time for 10 h followed by 90s switching time for 17h). The voltage was 6V/cm and the included angle 120°. Agarose gels were stained for 20 min with ethidium bromide in TBE 0.5X.

### Genetic instability assays

*LYS2* forward mutation assays were performed as described^66^. Independent colonies were inoculated into 1 mL YPD and grown non-selectively to saturation at 30°C. Cells were washed and resuspended in water prior to plating on SC medium (to determine the total number of viable cells) or alpha-aminoadipate medium (to select for *lys2^-^*mutants). Mutation rates were calculated using bz-rates^109^, based on measurements obtained from 7-8 independent colonies per fluctuation assays. At least four independent fluctuation assays were performed for each genotype.

Mutational profiling of the *LYS2* locus was achieved by deep sequencing of a *LYS2* amplicon generated from pool of *lys2^-^*mutants, based on a published procedure^110^. For each genotype, approximately *N*=90-100 *lys2^-^* independent mutants were pooled in equal proportions. Genomic DNA was extracted from each pool, as above, and the *LYS2* gene (CDS -381/+428 bp) was amplified by PCR using the Phusion enzyme and primers described in **Suppl. Table 4**. Full-length amplicons were purified, used for paired-end library preparation and sequenced using a NovaSeq X Plus platform (Novogene). Raw paired-end Illumina reads were processed using FastP, with a minimum quality score of 20, a maximum of 6% low-quality bases per read (-u 6), and a minimum read length of 30 bp, and further aligned to the *LYS2* amplicon using Bowtie2 (version 2.5.1). Aligned reads were filtered to retain only the reads mapped in a proper pair with a quality score ≥20 using Samtools (version 1.23) for subsequent SNP and INDELs detection. Variants were called using iVar (version 1.4.4), and based on their allele frequency. Sequencing depth was adjusted to ensure that mutations with allelic frequency between 0.65 x 1/*N* and 0.95 could be included^110^. The number of distinct mutations detected did not scale proportionally with the number of pooled mutants, consistent with the presence of mutational hotspots within the *LYS2* locus, as previously reported^66^. For each mutation, the estimated number of independent mutants carrying that alteration was thus calculated by multiplying the observed mutant allele frequency by the total number of pooled mutants. Structural variants affecting the *LYS2* locus, as well as the accuracy of mutational spectrum determination, were validated by PCR amplification and Sanger sequencing of a subset of independent *lys2^-^* clones. The chromosome XVI deletion assay was performed as follows. Independent colonies were inoculated into 1 mL YPD and grown non-selectively to saturation at 30°C. Cells were washed and resuspended in water prior to plating on SC medium (to determine the total number of viable cells) or 5-fluoroorotic acid medium (to select for *ura3^-^*mutants). Colonies growing on 5-FOA plates were replica-plated onto YPD medium supplemented with hygromycin B (Thermo Fisher Scientific, 300µg/mL) to identify *ura3^-^ HygR* colonies. Deletion frequencies were determined by dividing the number of *ura3^-^ HygR* mutants by the total number of viable cells. For each genotype, at least 4 independent replicates were performed. Each replicate represents the median value obtained from measurements performed on 6–8 independent colonies.

### Protein analyses

Yeast whole-cell extracts were obtained and analyzed by western blotting as previously reported^41^. The following validated antibodies were used: anti-Flag monoclonal antibody (M2, Sigma), 1:2000; anti-Dpm1 monoclonal antibody (Thermo Fisher Scientific), 1:1000; mouse peroxidase-conjugated antibodies (Jackson Immunoresearch). Membranes were revealed using chemiluminescent substrates, as above.

### Quantification and statistical analysis

Experiments were not randomized, and investigators were not blinded to group allocation during data collection or analysis. Sample sizes (n) were selected in accordance with standard practice in the field and correspond to the number of biological replicates (e.g., independent cultures). For fluctuation assays, each replicate (n≥4) represents the mutation rate or frequency determined using 6–8 independent measurements performed on individual colonies. Error bars correspond to standard deviations. The following statistical tests were used to evaluate the statistical differences between strains or proportions, as indicated: Two-tailed Mann-Whitney-Wilcoxon rank sum test (all Figures except **Fig. 1g-h**); Fisher-exact test (**Fig. 1g-h**). Box-plots were represented according to Tukey’s definition using Prism v10.6.1. Outliers were not represented in **Fig. 1f** & **Ext. Data Fig. 1f** for clarity but have been included in statistical analyses. The following convention was used for reporting statistical significance: * p<0.05; ** p<0.01; *** p<0.001; **** p<0.0001; ns, not significant. The complete NGS data generated during this study are available in the European Nucleotide Archive (ENA) database under accession number PRJEB121038.

## Acknowledgements

We are very grateful to Andres Aguilera for the *hpr1* mutant, and to Robert Crouch and Aziz El Hage for the *RNH201-RED* construct; to Nicolas Valentin (Imagoseine), Patricia Woni and Joel Marchand for valuable assistance with flow cytometry, media preparation, and computational resources, respectively; to Astrid Lancrey for technical help with MA line generation; to Pascale Lesage, Alain Nicolas, Rodney Rothstein and Vincent Vanoosthuyse for discussion and critical reading of the manuscript.

This work was supported by Agence Nationale pour la Recherche (ANR-18-CE12-0003, to B.P. and G.F.; ANR-21-CE12-0040, to B.P. and D.L.), Fondation ARC pour la recherche contre le Cancer (projet ARC, to B.P), Ligue Nationale contre le Cancer (Equipe Labellisée Ligue Contre le Cancer, to B.P., and PhD fellowship, to M.Z.), the BioSPC PhD program (to M.Z. and R.M.M.) and the Fondation Pour la Recherche Médicale (FDT202304016590, to R.M.M.). This work has received support from the investment program “France 2030” as part of the IdEx program (ANR-18-IDEX-0001) implemented by Université Paris Cité, under which the inIdEx project Formula is conducted.

## Author contributions

Conceptualization, M.Z., G.F., B.P.; Methodology, M.Z., F.P., S.O.D., N.A., D.L., B.P.; Investigation, all authors; Formal analysis, M.Z., F.P., S.O.D., N.A., U.A., B.P.; Writing - Original Draft, M.Z., B.P.; Writing - Review & Editing, all authors; Funding Acquisition, D.L., G.F., B.P.; Visualization, M.Z., B.P.; Supervision, F.P., D.L., G.F., B.P.; Project administration, B.P.

## References

1. Kellner, V. & Luke, B. Molecular and physiological consequences of faulty eukaryotic ribonucleotide excision repair. EMBO J. 39, e102309 (2020).

2. Brickner, J. R., Garzon, J. L. & Cimprich, K. A. Walking a tightrope: The complex balancing act of R-loops in genome stability. Mol. Cell 82, 2267–2297 (2022).

3. García-Muse, T. & Aguilera, A. R Loops: From Physiological to Pathological Roles. Cell 179, 604–618 (2019).

4. Balachander, S. et al. Ribonucleotide incorporation in yeast genomic DNA shows preference for cytosine and guanosine preceded by deoxyadenosine. Nat. Commun. 11, 2447 (2020).

5. Clausen, A. R. et al. Tracking replication enzymology in vivo by genome-wide mapping of ribonucleotide incorporation. Nat. Struct. Mol. Biol. 22, 185–191 (2015).

6. Chédin, F., Hartono, S. R., Sanz, L. A. & Vanoosthuyse, V. Best practices for the visualization, mapping, and manipulation of R-loops. EMBO J. 40, e106394 (2021).

7. Nick McElhinny, S. A., et al. Abundant ribonucleotide incorporation into DNA by yeast replicative polymerases. Proc. Natl. Acad. Sci. U. S. A. 107, 4949–4954 (2010).

8. Williams, J. S., Lujan, S. A. & Kunkel, T. A. Processing ribonucleotides incorporated during eukaryotic DNA replication. Nat. Rev. Mol. Cell Biol. 17, 350–363 (2016).

9. Wahba, L., Costantino, L., Tan, F. J., Zimmer, A. & Koshland, D. S1-DRIP-seq identifies high expression and polyA tracts as major contributors to R-loop formation. Genes Dev 30, 1327–38 (2016).

10. Sanz, L. A. et al. Prevalent, Dynamic, and Conserved R-Loop Structures Associate with Specific Epigenomic Signatures in Mammals. Mol. Cell 63, 167–178 (2016).

11. Bonnet, A. et al. Introns Protect Eukaryotic Genomes from Transcription-Associated Genetic Instability. Mol. Cell 67, 608–621.e6 (2017).

12. Luna, R., Rondón, A. G., Pérez-Calero, C., Salas-Armenteros, I. & Aguilera, A. The THO Complex as a Paradigm for the Prevention of Cotranscriptional R-Loops. Cold Spring Harb. Symp. Quant. Biol. 84, 105–114 (2019).

13. Cerritelli, S. M. & Crouch, R. J. RNases H: Multiple roles in maintaining genome integrity. DNA Repair 84, 102742 (2019).

14. Richard, P. & Manley, J. L. R Loops and Links to Human Disease. J Mol Biol 429, 3168–3180 (2017).

15. Wells, J. P., White, J. & Stirling, P. C. R Loops and Their Composite Cancer Connections. Trends Cancer 5, 619–631 (2019).

16. Aiello, U. et al. Sen1 is a key regulator of transcription-driven conflicts. Mol. Cell 82, 2952–2966.e6 (2022).

17. Arana, M. E. et al. Transcriptional responses to loss of RNase H2 in Saccharomyces cerevisiae. DNA Repair 11, 933–941 (2012).

18. Han, Z. et al. A role for human senataxin in contending with pausing and backtracking during transcript elongation. Mol. Cell 85, 4166–4182.e10 (2025).

19. Gomez-Gonzalez, B. & Aguilera, A. Activation-induced cytidine deaminase action is strongly stimulated by mutations of the THO complex. Proc Natl Acad Sci U A 104, 8409–14 (2007).

20. Kim, N. et al. Mutagenic processing of ribonucleotides in DNA by yeast topoisomerase I. Science 332, 1561–1564 (2011).

21. Makharashvili, N. et al. Sae2/CtIP prevents R-loop accumulation in eukaryotic cells. eLife 7, (2018).

22. Sollier, J. et al. Transcription-coupled nucleotide excision repair factors promote R-loop-induced genome instability. Mol Cell 56, 777–85 (2014).

23. Sparks, J. L. & Burgers, P. M. Error-free and mutagenic processing of topoisomerase 1-provoked damage at genomic ribonucleotides. EMBO J. 34, 1259–1269 (2015).

24. Su, X. A. & Freudenreich, C. H. Cytosine deamination and base excision repair cause R-loop-induced CAG repeat fragility and instability in Saccharomyces cerevisiae. Proc Natl Acad Sci U A 114, E8392–E8401 (2017).

25. Williams, J. S. et al. Topoisomerase 1-mediated removal of ribonucleotides from nascent leading-strand DNA. Mol. Cell 49, 1010–1015 (2013).

26. Hamperl, S., Bocek, M. J., Saldivar, J. C., Swigut, T. & Cimprich, K. A. Transcription-Replication Conflict Orientation Modulates R-Loop Levels and Activates Distinct DNA Damage Responses. Cell 170, 774–786.e19 (2017).

27. Lang, K. S. et al. Replication-Transcription Conflicts Generate R-Loops that Orchestrate Bacterial Stress Survival and Pathogenesis. Cell 170, 787–799.e18 (2017).

28. Lazzaro, F. et al. RNase H and postreplication repair protect cells from ribonucleotides incorporated in DNA. Mol. Cell 45, 99–110 (2012).

29. Costantino, L. & Koshland, D. Genome-wide Map of R-Loop-Induced Damage Reveals How a Subset of R-Loops Contributes to Genomic Instability. Mol. Cell 71, 487–497.e3 (2018).

30. Aguilera, A. & Klein, H. L. Genetic control of intrachromosomal recombination in Saccharomyces cerevisiae. I. Isolation and genetic characterization of hyper-recombination mutations. Genetics 119, 779–790 (1988).

31. Nick McElhinny, S. A., et al. Genome instability due to ribonucleotide incorporation into DNA. Nat. Chem. Biol. 6, 774–781 (2010).

32. Cornelio, D. A., Sedam, H. N. C., Ferrarezi, J. A., Sampaio, N. M. V. & Argueso, J. L. Both R-loop removal and ribonucleotide excision repair activities of RNase H2 contribute substantially to chromosome stability. DNA Repair 52, 110–114 (2017).

33. Huertas, P. & Aguilera, A. Cotranscriptionally formed DNA:RNA hybrids mediate transcription elongation impairment and transcription-associated recombination. Mol Cell 12, 711–21 (2003).

34. Wahba, L., Amon, J. D., Koshland, D. & Vuica-Ross, M. RNase H and multiple RNA biogenesis factors cooperate to prevent RNA:DNA hybrids from generating genome instability. Mol Cell 44, 978–88 (2011).

35. Santos-Rosa, H. & Aguilera, A. Increase in incidence of chromosome instability and non-conservative recombination between repeats in Saccharomyces cerevisiae hpr1 delta strains. Mol. Gen. Genet. MGG 245, 224–236 (1994).

36. Amon, J. D. & Koshland, D. RNase H enables efficient repair of R-loop induced DNA damage. eLife 5, e20533 (2016).

37. Schroeder, J. W. et al. RNase H genes cause distinct impacts on RNA:DNA hybrid formation and mutagenesis genome wide. Sci. Adv. 9, eadi5945 (2023).

38. Promonet, A. et al. Topoisomerase 1 prevents replication stress at R-loop-enriched transcription termination sites. Nat. Commun. 11, 3940 (2020).

39. Heuzé, J. et al. RNase H2 degrades toxic RNA:DNA hybrids behind stalled forks to promote replication restart. EMBO J. 42, e113104 (2023).

40. Stoy, H. et al. Direct visualization of transcription-replication conflicts reveals post-replicative DNA:RNA hybrids. Nat. Struct. Mol. Biol. 30, 348–359 (2023).

41. Mangione, R. M. et al. DNA lesions can frequently precede DNA:RNA hybrid accumulation. Nat. Commun. 16, 2401 (2025).

42. Saur, F. et al. Transcriptional repression facilitates RNA:DNA hybrid accumulation at DNA double-strand breaks. Nat. Cell Biol. 27, 992–1005 (2025).

43. Pessina, F. et al. Functional transcription promoters at DNA double-strand breaks mediate RNA-driven phase separation of damage-response factors. Nat. Cell Biol. 21, 1286–1299 (2019).

44. Niehrs, C. & Luke, B. Regulatory R-loops as facilitators of gene expression and genome stability. Nat. Rev. Mol. Cell Biol. 21, 167–178 (2020).

45. Liu, L., Sun, D. & Liu, H. Comprehensive Mutational Landscape of Yeast Mutator Strains Reveals the Genetic Basis of Mutational Signatures in Cancer. Mol. Biol. Evol. 42, msaf252 (2025).

46. Loeillet, S. et al. Trajectory and uniqueness of mutational signatures in yeast mutators. Proc. Natl. Acad. Sci. U. S. A. 117, 24947–24956 (2020).

47. Crossley, M. P., Bocek, M. J., Hamperl, S., Swigut, T. & Cimprich, K. A. qDRIP: a method to quantitatively assess RNA-DNA hybrid formation genome-wide. Nucleic Acids Res. 48, e84 (2020).

48. Wahba, L., Costantino, L., Tan, F. J., Zimmer, A. & Koshland, D. S1-DRIP-seq identifies high expression and polyA tracts as major contributors to R-loop formation. Genes Dev. 30, 1327–1338 (2016).

49. Penzo, A. et al. A R-loop sensing pathway mediates the relocation of transcribed genes to nuclear pore complexes. Nat. Commun. 14, 5606 (2023).

50. Caudal, É. et al. Pan-transcriptome reveals a large accessory genome contribution to gene expression variation in yeast. Nat. Genet. 56, 1278–1287 (2024).

51. Sharp, N. P., Sandell, L., James, C. G. & Otto, S. P. The genome-wide rate and spectrum of spontaneous mutations differ between haploid and diploid yeast. Proc. Natl. Acad. Sci. U. S. A. 115, E5046–E5055 (2018).

52. Serero, A., Jubin, C., Loeillet, S., Legoix-Ne, P. & Nicolas, A. G. Mutational landscape of yeast mutator strains. Proc Natl Acad Sci U A 111, 1897–902 (2014).

53. Lynch, M. et al. A genome-wide view of the spectrum of spontaneous mutations in yeast. Proc. Natl. Acad. Sci. U. S. A. 105, 9272–9277 (2008).

54. Harari, Y., Ram, Y., Rappoport, N., Hadany, L. & Kupiec, M. Spontaneous Changes in Ploidy Are Common in Yeast. Curr. Biol. 28, 825–835.e4 (2018).

55. Venkataram, S. et al. Development of a Comprehensive Genotype-to-Fitness Map of Adaptation-Driving Mutations in Yeast. Cell 166, 1585–1596.e22 (2016).

56. Williams, J. S. et al. Genome-wide mutagenesis resulting from topoisomerase 1-processing of unrepaired ribonucleotides in DNA. DNA Repair 84, 102641 (2019).

57. Chon, H. et al. RNase H2 roles in genome integrity revealed by unlinking its activities. Nucleic Acids Res. 41, 3130–3143 (2013).

58. Yang, X. et al. RNA-DNA hybrids regulate meiotic recombination. Cell Rep. 37, 110097 (2021).

59. Ghodgaonkar, M. M. et al. Ribonucleotides Misincorporated into DNA Act as Strand-Discrimination Signals in Eukaryotic Mismatch Repair. Mol. Cell 50, 323–332 (2013).

60. Lujan, S. A., Williams, J. S., Clausen, A. R., Clark, A. B. & Kunkel, T. A. Ribonucleotides Are Signals for Mismatch Repair of Leading-Strand Replication Errors. Mol. Cell 50, 437–443 (2013).

61. McVey, M., Khodaverdian, V. Y., Meyer, D., Cerqueira, P. G. & Heyer, W.-D. Eukaryotic DNA Polymerases in Homologous Recombination. Annu. Rev. Genet. 50, 393–421 (2016).

62. Curcio, M. J., Lutz, S. & Lesage, P. The Ty1 LTR-retrotransposon of budding yeast, Saccharomyces cerevisiae. Microbiol. Spectr. 3, 1–35 (2015).

63. Hoang, M. L. et al. Competitive repair by naturally dispersed repetitive DNA during non-allelic homologous recombination. PLoS Genet. 6, e1001228 (2010).

64. Brown, R. E. et al. The RNA export and RNA decay complexes THO and TRAMP prevent transcription-replication conflicts, DNA breaks, and CAG repeat contractions. PLoS Biol. 20, e3001940 (2022).

65. Mérida-Cerro, J. A., Maraver-Cárdenas, P., Rondón, A. G. & Aguilera, A. Rat1 promotes premature transcription termination at R-loops. Nucleic Acids Res. gkae033 (2024) doi:10.1093/nar/gkae033.

66. Lippert, M. J., Freedman, J. A., Barber, M. A. & Jinks-Robertson, S. Identification of a distinctive mutation spectrum associated with high levels of transcription in yeast. Mol. Cell. Biol. 24, 4801–4809 (2004).

67. Flores-Rozas, H. & Kolodner, R. D. The Saccharomyces cerevisiae MLH3 gene functions in MSH3-dependent suppression of frameshift mutations. Proc. Natl. Acad. Sci. U. S. A. 95, 12404–12409 (1998).

68. Kunz, B. A., Ramachandran, K. & Vonarx, E. J. DNA sequence analysis of spontaneous mutagenesis in Saccharomyces cerevisiae. Genetics 148, 1491–1505 (1998).

69. Senoussi, I. et al. Mechanism of trinucleotide repeat expansion by MutSβ-MutLγ and contraction by FAN1. Nat. Commun. 16, 9445 (2025).

70. El Hage, A., Webb, S., Kerr, A. & Tollervey, D. Genome-wide distribution of RNA-DNA hybrids identifies RNase H targets in tRNA genes, retrotransposons and mitochondria. PLoS Genet 10, e1004716 (2014).

71. Winston, F., Durbin, K. J. & Fink, G. R. The SPT3 gene is required for normal transcription of Ty elements in S. cerevisiae. Cell 39, 675–682 (1984).

72. Prado, F. & Aguilera, A. Impairment of replication fork progression mediates RNA polII transcription-associated recombination. EMBO J. 24, 1267–1276 (2005).

73. McGuffee, S. R., Smith, D. J. & Whitehouse, I. Quantitative, genome-wide analysis of eukaryotic replication initiation and termination. Mol. Cell 50, 123–135 (2013).

74. Chang, E. Y.-C. et al. RECQ-like helicases Sgs1 and BLM regulate R-loop-associated genome instability. J. Cell Biol. 216, 3991–4005 (2017).

75. Schroeder, J. W., Randall, J. R., Hirst, W. G., O’Donnell, M. E. & Simmons, L. A. Mutagenic cost of ribonucleotides in bacterial DNA. Proc. Natl. Acad. Sci. U. S. A. 114, 11733–11738 (2017).

76. Conover, H. N. et al. Stimulation of Chromosomal Rearrangements by Ribonucleotides. Genetics 201, 951–961 (2015).

77. O’Connell, K., Jinks-Robertson, S. & Petes, T. D. Elevated Genome-Wide Instability in Yeast Mutants Lacking RNase H Activity. Genetics 201, 963–975 (2015).

78. Hoyt, S. J. et al. From telomere to telomere: The transcriptional and epigenetic state of human repeat elements. Science 376, eabk3112 (2022).

79. O’Donnell, S. et al. Telomere-to-telomere assemblies of 142 strains characterize the genome structural landscape in Saccharomyces cerevisiae. Nat. Genet. 55, 1390–1399 (2023).

80. Chan, J. E. & Kolodner, R. D. A genetic and structural study of genome rearrangements mediated by high copy repeat Ty1 elements. PLoS Genet. 7, e1002089 (2011).

81. Groh, M., Lufino, M. M., Wade-Martins, R. & Gromak, N. R-loops associated with triplet repeat expansions promote gene silencing in Friedreich ataxia and fragile X syndrome. PLoS Genet 10, e1004318 (2014).

82. Lin, Y., Dent, S. Y. R., Wilson, J. H., Wells, R. D. & Napierala, M. R loops stimulate genetic instability of CTG·CAG repeats. Proc. Natl. Acad. Sci. 107, 692–697 (2010).

83. Neil, A. J., Liang, M. U., Khristich, A. N., Shah, K. A. & Mirkin, S. M. RNA-DNA hybrids promote the expansion of Friedreich’s ataxia (GAA)n repeats via break-induced replication. Nucleic Acids Res. 46, 3487–3497 (2018).

84. Reddy, K. et al. Processing of double-R-loops in (CAG)·(CTG) and C9orf72 (GGGGCC)·(GGCCCC) repeats causes instability. Nucleic Acids Res. 42, 10473–10487 (2014).

85. Freudenreich, C. H. R-loops: targets for nuclease cleavage and repeat instability. Curr. Genet. 64, 789–794 (2018).

86. Kadyrova, L. Y., Gujar, V., Burdett, V., Modrich, P. L. & Kadyrov, F. A. Human MutLγ, the MLH1-MLH3 heterodimer, is an endonuclease that promotes DNA expansion. Proc. Natl. Acad. Sci. U. S. A. 117, 3535–3542 (2020).

87. Lin, Y., Dion, V. & Wilson, J. H. Transcription promotes contraction of CAG repeat tracts in human cells. Nat. Struct. Mol. Biol. 13, 179–180 (2006).

88. Reijns, M. A. M. et al. Signatures of TOP1 transcription-associated mutagenesis in cancer and germline. Nature 602, 623–631 (2022).

89. Bayona-Feliu, A. et al. The chromatin network helps prevent cancer-associated mutagenesis at transcription-replication conflicts. Nat. Commun. 14, 6890 (2023).

90. Houlahan, K. E. et al. Complex rearrangements fuel ER+ and HER2+ breast tumours. Nature 638, 510–518 (2025).

91. Jalan, M. et al. RAD52 resolves transcription-replication conflicts to mitigate R-loop induced genome instability. Nat. Commun. 15, 7776 (2024).

92. Ellrott, K. et al. Scalable Open Science Approach for Mutation Calling of Tumor Exomes Using Multiple Genomic Pipelines. Cell Syst. 6, 271–281.e7 (2018).

93. Sondka, Z. et al. COSMIC: a curated database of somatic variants and clinical data for cancer. Nucleic Acids Res. 52, D1210–D1217 (2024).

94. Sikorski, R. S. & Boeke, J. D. In vitro mutagenesis and plasmid shuffling: from cloned gene to mutant yeast. Methods Enzymol. 194, 302–318 (1991).

95. Garcia-Rubio, M. et al. Different physiological relevance of yeast THO/TREX subunits in gene expression and genome integrity. Mol Genet Genomics 279, 123–32 (2008).

96. Hu, B. et al. Biological chromodynamics: a general method for measuring protein occupancy across the genome by calibrating ChIP-seq. Nucleic Acids Res. 43, e132 (2015).

97. Ramírez, F. et al. deepTools2: a next generation web server for deep-sequencing data analysis. Nucleic Acids Res. 44, W160–W165 (2016).

98. Xu, Z. et al. Bidirectional promoters generate pervasive transcription in yeast. Nature 457, 1033–7 (2009).

99. Nick McElhinny, S. A., et al. Genome instability due to ribonucleotide incorporation into DNA. Nat. Chem. Biol. 6, 774–781 (2010).

100. Cingolani, P. et al. A program for annotating and predicting the effects of single nucleotide polymorphisms, SnpEff: SNPs in the genome of Drosophila melanogaster strain w1118; iso-2; iso-3. Fly (Austin) 6, 80–92 (2012).

101. Denis, E. et al. Extracting high molecular weight genomic DNA from Saccharomyces cerevisiae. Protoc. Exch. 10.1038/protex.2018.076 (2018) doi:10.1038/protex.2018.076.

102. Koren, S. et al. Canu: scalable and accurate long-read assembly via adaptive k-mer weighting and repeat separation. Genome Res. 27, 722–736 (2017).

103. Liu, H., Wu, S., Li, A. & Ruan, J. SMARTdenovo: a de novo assembler using long noisy reads. GigaByte 2021, gigabyte15 (2021).

104. Hu, J. et al. NextDenovo: an efficient error correction and accurate assembly tool for noisy long reads. Genome Biol. 25, 107 (2024).

105. Vaser, R., Sović, I., Nagarajan, N. & Šikić, M. Fast and accurate de novo genome assembly from long uncorrected reads. Genome Res. 27, 737–746 (2017).

106. Kolmogorov, M., Raney, B., Paten, B. & Pham, S. Ragout-a reference-assisted assembly tool for bacterial genomes. Bioinformatics 30, i302–309 (2014).

107. Krzywinski, M. et al. Circos: an information aesthetic for comparative genomics. Genome Res. 19, 1639–1645 (2009).

108. Török, T., Rockhold, D. & King, A. D. Use of electrophoretic karyotyping and DNA-DNA hybridization in yeast identification. Int. J. Food Microbiol. 19, 63–80 (1993).

109. Gillet-Markowska, A., Louvel, G. & Fischer, G. bz-rates: A Web Tool to Estimate Mutation Rates from Fluctuation Analysis. G3 5, 2323–2327 (2015).

110. Jiang, P. et al. A modified fluctuation assay reveals a natural mutator phenotype that drives mutation spectrum variation within Saccharomyces cerevisiae. eLife 10, e68285 (2021).

