## Supplementary Information for "DNA:RNA hybrid mutational landscapes reveal a common route to genetic instability in evolving genomes"

Includes

**Ext. Data Figures 1-6**

**Suppl. Table 1 – List of genetic alterations detected in all MA line analyses** (*provided as an independent Excel spreadsheet*)

**Suppl. Table 2 - Yeast strains used in this study**

**Suppl. Table 3 - Plasmids used in this study**

**Suppl. Table 4 - Oligonucleotides used in this study**

**Supplementary references**

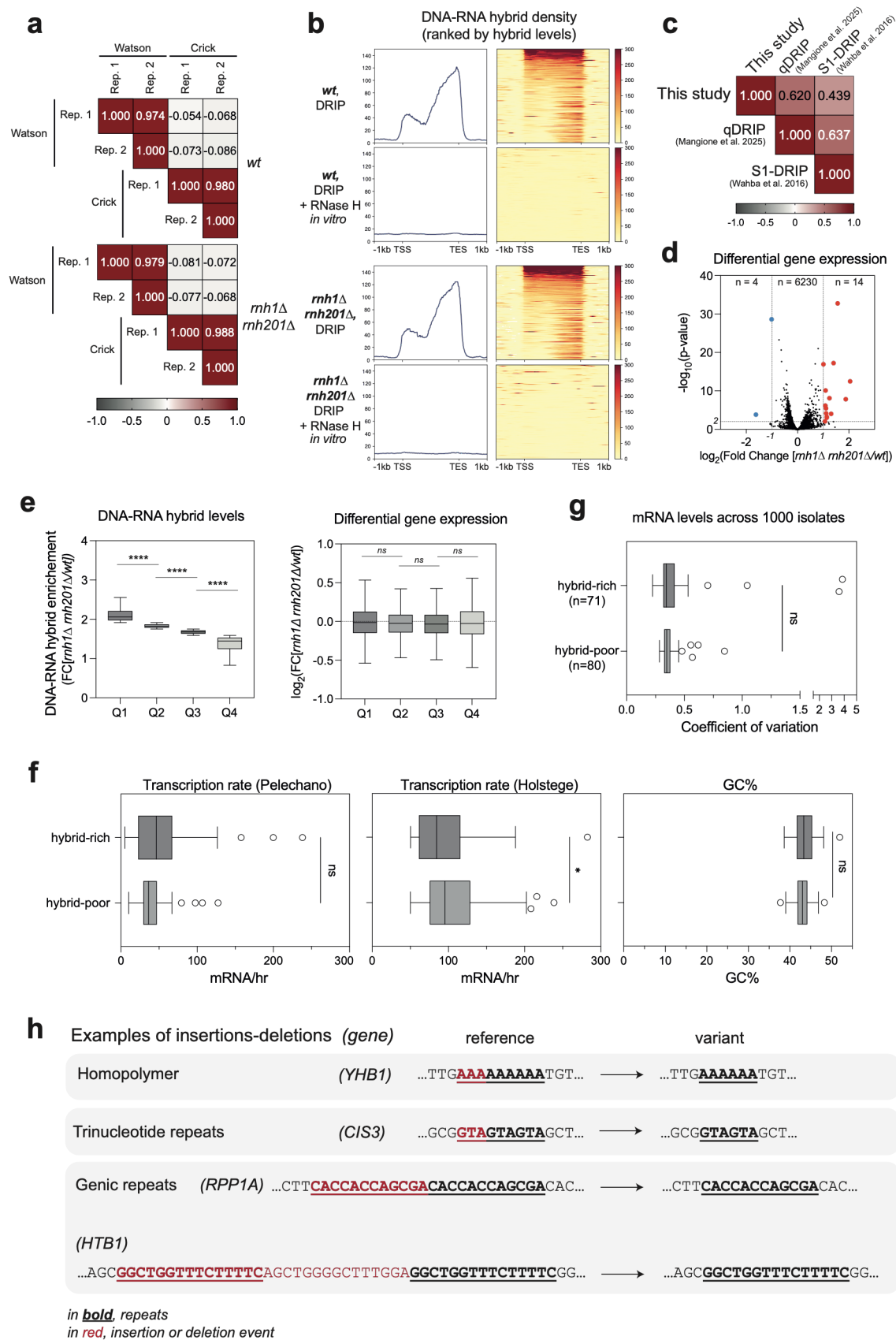

Ext. Data Figure 1, related to Fig. 1. Characterization of DNA:RNA hybrid maps for intronless and intron-containing gene sets. See legend on next page.

**Ext. Data Figure 1, related to Fig. 1. Characterization of DNA:RNA hybrid maps for intronless and intron-containing gene sets.** **a**, Correlation analysis for DRIP-seq biological replicates (Rep. 1 and Rep. 2), comparing signal from Watson or Crick strands. For each comparison, the Pearson correlation coefficient is indicated. **b**, Metagene and heatmap analysis of DNA:RNA hybrid levels (DRIP signals in immunoprecipitates) at highly-transcribed genes (n=151) aligned at their Transcription Start Site (TSS) and Transcription End Site (TES). When indicated, DNA samples were treated with RNase H *in vitro* prior to immunoprecipitation. Note that spike-in calibration was not introduced in this specific analysis since RNase H also degrades the spike-in hybrids. **c**, Correlation analysis for distinct DNA:RNA hybrid datasets obtained from control cells: qDRIP (this study ; Mangione et al., 2025<sup>1</sup>); S1-DRIP-seq (*wt* cells, Wahba et al.<sup>2</sup>). **d**, RNA-seq analysis of *wt* and *rnh1Δ rnh201Δ* cells. Differential expression ($\log_2$ Fold Change [*rnh1Δ rnh201Δ/wt*]) and associated significance ($-\log_{10}$ p-value, based on three replicates) are represented for the whole transcriptome. Transcripts that are up-regulated ($\log_2$ FC > 1; $p < 0.05$) or down-regulated ($\log_2$ FC < -1; $p < 0.05$) upon inactivation of RNases H appear in red and blue, respectively. **e**, Genes were splitted into quartiles based on their enrichment in hybrids in the *rnh1Δ rnh201Δ* mutant relative to *wt* (*left panel*). Differential gene expression (from transcriptome data, $\log_2$ Fold Change [*rnh1Δ rnh201Δ/wt*]) is represented for each of these quartiles (*right panel*). **f**, *Left and mid panel*, Transcription rates (mRNA/hour) for hybrid-prone (intronless) and hybrid-poor (intron-containing) gene groups, using measurements from Pelechano et al.<sup>3</sup> (*left panel*) and Holstege et al.<sup>4</sup> (*mid panel*). *Right panel*, GC content (%) for hybrid-prone (intronless) and hybrid-poor (intron-containing) gene groups (excluding intronic sequences). **g**, For each gene in both groups, transcriptomic data from 969 distinct *S. cerevisiae* isolates were retrieved, and the coefficient of variation of mRNA abundance across the 969 transcriptomes was calculated. The distributions of coefficients of variation for the hybrid-rich (n=71) and hybrid-poor (n=80) gene groups were then represented as box plots. **h**, Reference and variant sequences are shown for examples of intronless, hybrid-rich genes in which INDELs are observed within repetitive elements: homopolymers, trinucleotide repeats or genic repeats. \* $p < 0.05$; \*\*\*\* $p < 0.0001$; ns, not significant (Mann-Whitney-Wilcoxon rank sum test).

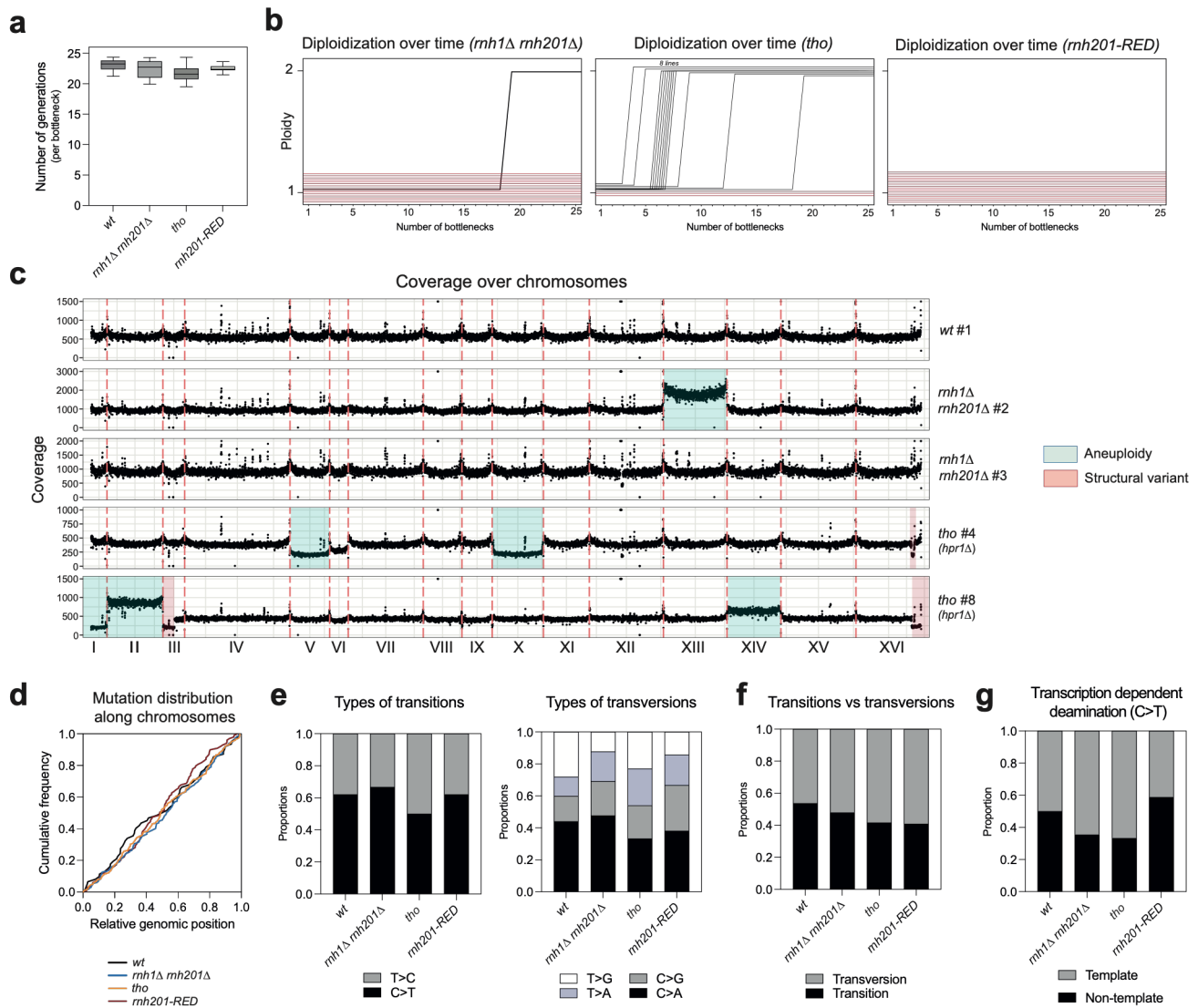

**Ext. Data Figure 2, related to Fig. 2. Phenotypes and mutational spectra of mutation accumulation lines.** **a**, Number of generations per bottleneck for the different genotypes assessed in MA lines analyses (*wt*, $n=16$; *rnh1Δ rnh201Δ*, $n=15$; *tho* (*hpr1Δ*), $n=16$; *rnh201-RED*, $n=16$). **b**, Timeline of diploidization for *rnh1Δ rnh201Δ* (left panel), *tho* (*hpr1Δ*, central panel), and *rnh201-RED* (right panel) MA lines. Each line represents an individual MA line (red, remaining haploid; black, diploidized at the indicated bottleneck). **c**, Example of whole-genome coverage for 5 representative MA lines: *wt* (line #1), *rnh1Δ rnh201Δ* (lines #2 and #3) and *tho* (*hpr1Δ*, lines #4 and #8). Positions of typical aneuploidies and structural variants are highlighted. **d**, Distribution of genetic alterations along chromosomes. The cumulative frequency of SNVs and INDELs is plotted as a function of normalized chromosomal position. **e-g**, Types of SNVs in the different MA lines analyzed (*wt*; *rnh1Δ rnh201Δ*; *tho* (*hpr1Δ*); *rnh201-RED*).

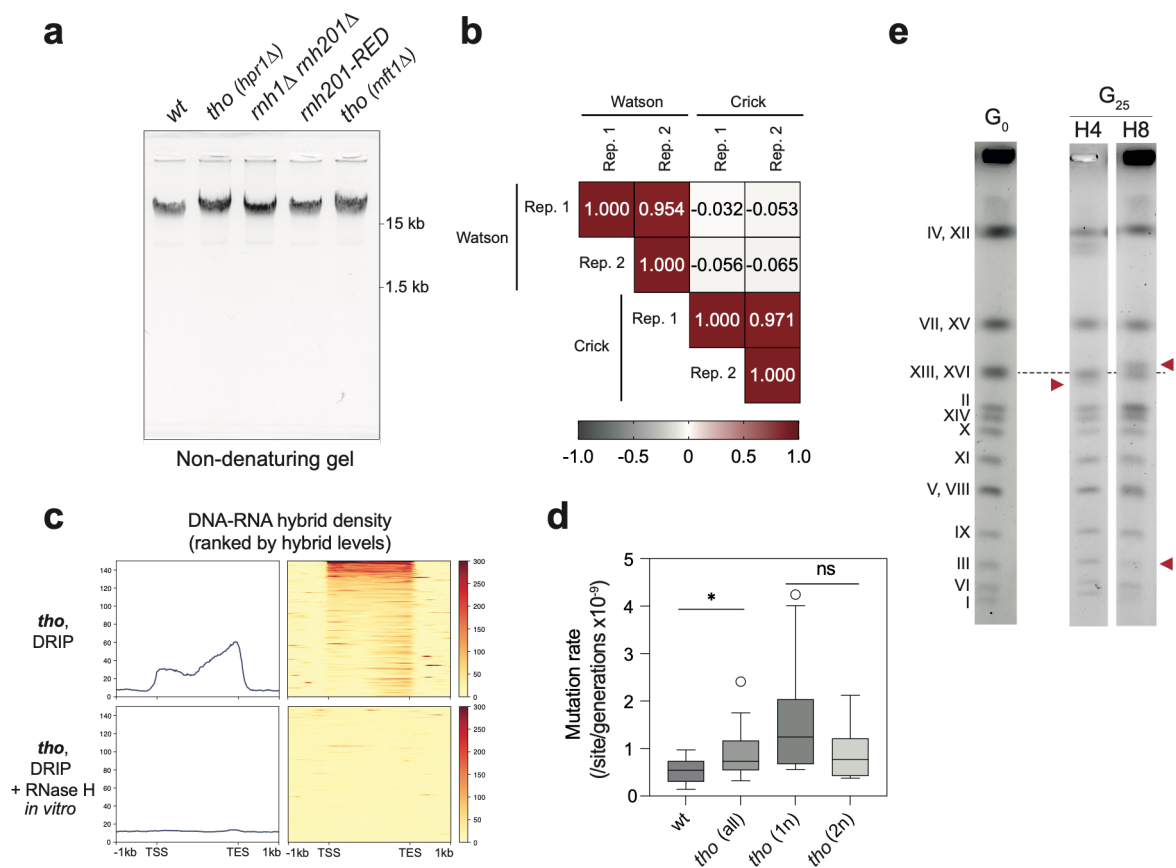

**Ext. Data Figure 3, related to Fig. 3. DNA:RNA hybrid phenotypes and structural variants in *tho* and *rnh201-RED* lines.** **a**, Non-denaturing agarose electrophoresis analysis of KCl-treated genomic DNA samples from the indicated strains. These samples correspond to those shown in **Fig. 3b**, in which they were treated with KOH and analyzed under alkaline electrophoresis conditions. The positions of the molecular weight markers are indicated (kb). **b**, Correlation analysis for DRIP-seq biological replicates (*tho* Rep. 1 and *tho* Rep. 2), comparing signal from Watson or Crick strands. For each comparison, the Pearson correlation coefficient is indicated. **c**, Metagene and heatmap analysis of DNA:RNA hybrid levels in *tho* (*hpr1*Δ) cells (DRIP signals in immunoprecipitates) at highly-transcribed genes (n=151) aligned at their TSS and TES. When indicated, DNA samples were treated with RNase H *in vitro* prior to immunoprecipitation. Calibration was not introduced in this analysis since RNase H also degrades the spike-in hybrids. **d**, Mutation rates for *tho* (*hpr1*Δ) lines according to their ploidy state. Mutations that were heterozygous in diploid lines were assumed to have arisen after diploidization, whereas homozygous mutations were assumed to have arisen before the diploidization event. Haploid mutation rates (1n) were estimated by dividing the number of

homozygous mutations by the haploid genome size and the number of generations spent in the haploid state (from **Ext. Data Fig. 2b**). Diploid mutation rates ($2n$) were estimated by dividing the number of heterozygous mutations by the diploid genome size and the number of generations spent in the diploid state. **e**, Representative PFGE analysis of genomic DNA from MA lines displaying structural variants. The chromosomal migration pattern of the parental $G_0$ isolate is shown, together with those of two $G_{25}$ *hpr1Δ* lines exhibiting SV (H4 and H8). Red arrows point to structural variants (see the interpretation below, **Ext. Data Fig. 4b-c**). \* $p < 0.05$; ns, not significant (Mann-Whitney-Wilcoxon rank sum test).

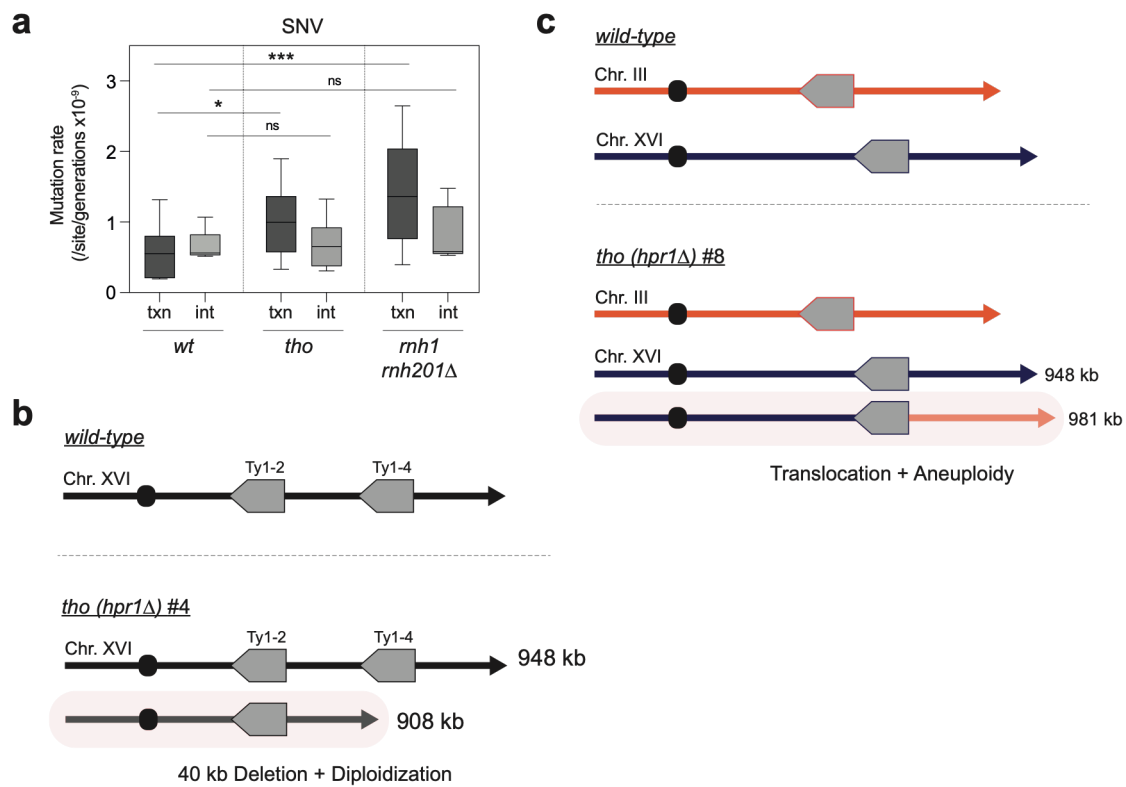

**Ext. Data Figure 4, related to Fig. 4. Repeat-associated genetic alterations in mutation accumulation lines.** **a**, Mutation rates (SNVs) were calculated separately for intergenic (*int*) and transcribed (*txn*) regions using previously published transcript annotations<sup>5</sup>. **b**, Example of a heterozygous deletion scored within chromosome XVI in the *tho (hpr1Δ)* MA line #4. **c**, Example of a non-reciprocal translocation scored between chromosomes XVI and III in the *tho (hpr1Δ)* MA line #8. Note that both structural variants exhibit breakpoints at Ty1 elements. \* $p < 0.05$; \*\*\* $p < 0.001$; ns, not significant (Mann-Whitney-Wilcoxon rank sum test).

**a** Examples of *LYS2* rearrangements

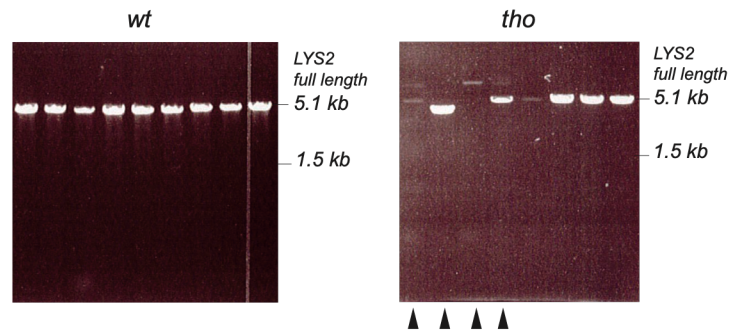

**b** Examples of INDELs identified by NGS

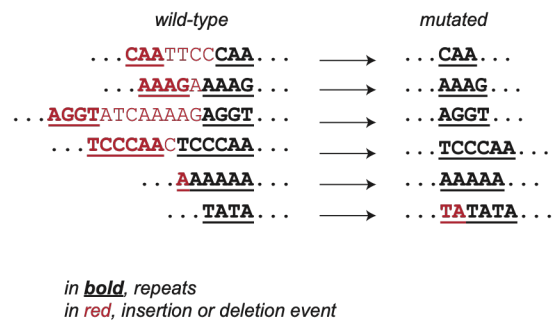

**Ext. Data Figure 5, related to Fig. 5. Examples of genetic alterations scored at the *LYS2* locus.**

**a**, Representative examples of *LYS2* amplicons generated from individual *lys2* mutants arising in the *wt* (left panel) or *tho* (*mft1Δ*; right panel) backgrounds. Note that a subset of *tho* mutants exhibit rearrangements affecting the *LYS2* locus (arrows), consistent with a previous study similarly analyzing a *URA3* reporter<sup>6</sup>. These rearrangements were classified as structural variants in the mutational spectra presented in **Fig. 5j**. **b**, Examples of INDELs scored within the *LYS2* locus in *tho* mutants, as identified by NGS.

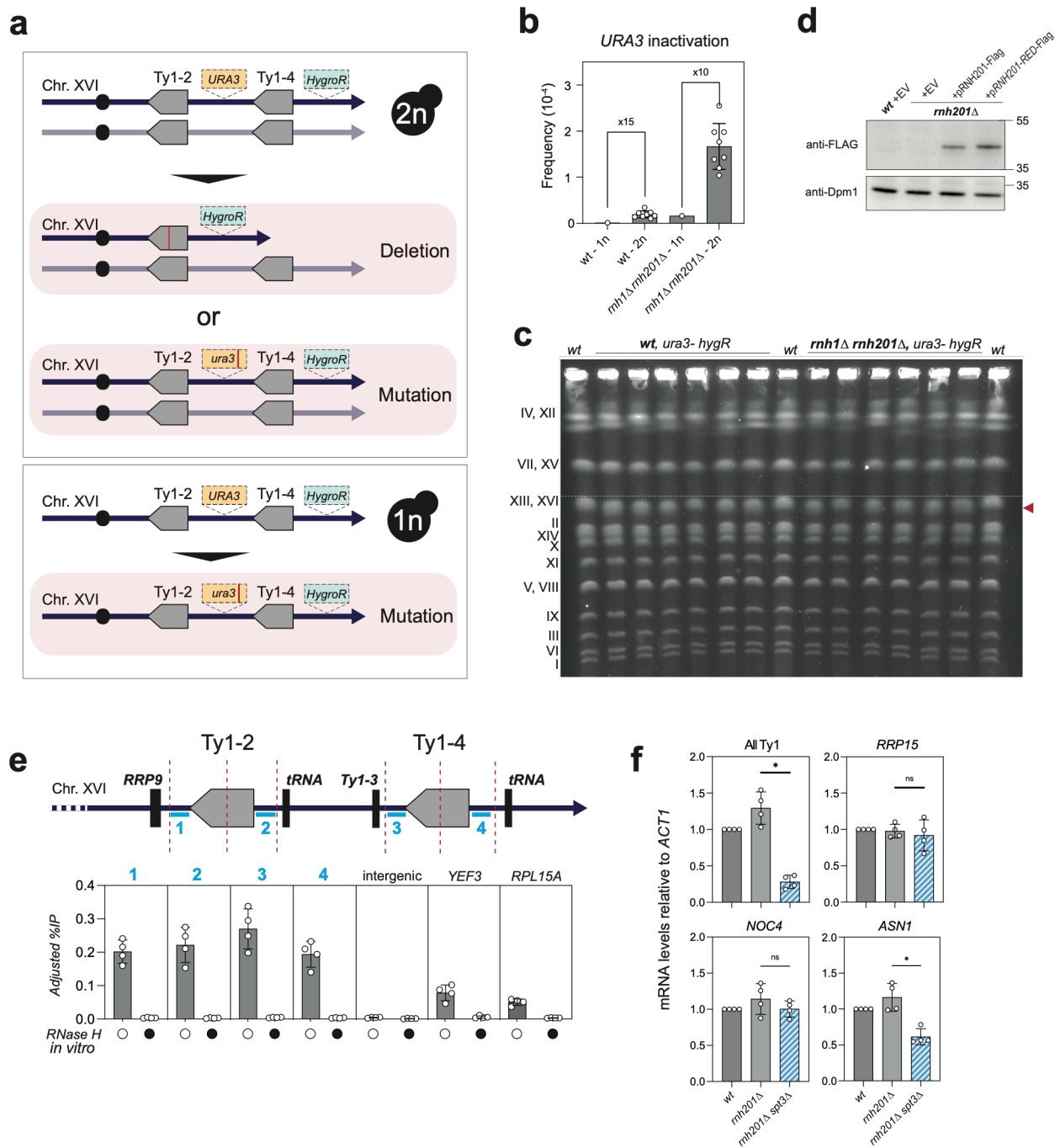

Ext. Data Figure 6, related to Fig. 6. Control experiments for the deletion reporter assay.

See legend on next page.

**Ext. Data Figure 6, related to Fig. 6. Control experiments for the deletion reporter assay.**

**a**, Schematic representation of the origin of *ura3* mutants in diploid vs. haploid strains carrying the deletion reporter. The red bar within the *URA3* locus represents an inactivating mutation. **b**, *URA3* inactivation frequency in *wt* and *rnh1Δ rnh201Δ* strains, either haploid (1n) or diploid (2n). **c**, PFGE analysis of diploid *ura3<sup>-</sup> hygR* colonies isolated in the deletion reporter assay. Note the occurrence of a faster-migrating version of chromosome XVI (red arrow), compatible with the 40 kb deletion, in *ura3<sup>-</sup> hygR* isolates, either *wt* (n=6) or *rnh1Δ rnh201Δ* (n=6). The position of chromosome XVI in *wt* parental controls is indicated by a dotted line. **d**, Expression levels of Rnh201, either *wt* or RED (*P45D Y219A*), as expressed in the indicated strains, and detected by anti-Flag western blotting. Dpm1 was used as a loading control. The position of molecular weights is indicated (kDa). EV, empty vector. **e**, DNA:RNA hybrid detection (DRIP-qPCR; adjusted % of IP; mean±SD; n=4) at chromosome XVI Ty1-2 and Ty1-4 loci in *wt* cells. The positions of restriction fragments (delineated by red dotted lines) and PCR amplicons (blue bars) are represented. When indicated, DNA extracts were treated with RNase H *in vitro* prior to immunoprecipitation (black circles). **f**, RT-qPCR quantification of mRNA levels from the indicated loci in the indicated strains. Values were normalized to *ACT1* RNA levels and expressed relative to *wt* (mean±SD, n=4). *RRP15*, *NOC4* and *ASN1* are three protein-coding genes located within the ~40-kb interval separating the Ty1-2 and Ty1-4 loci on chromosome XVI. Note that the primers used for Ty1 RNA detection do not discriminate among RNAs transcribed from distinct Ty1 loci and therefore measure total Ty1 transcript levels. \* p<0.05; ns, not significant (Mann-Whitney-Wilcoxon rank sum test).

**Suppl. Table 2 - Yeast strains used in this study**

| Code | Name | Relevant genotype | Source |
| --- | --- | --- | --- |
| ***S. cerevisiae*** |  |  |  |
| BY4741 | *wt* | *MATa ura3 his3 leu2 met15 LYS2* | *Euroscarf* |
| BY4742 | *wt* | *MATalpha ura3 his3 leu2 MET15 lys2* | *Euroscarf* |
| yBP2213 | *wt (G<sub>0</sub>)* | *(BY4741)* | *This study (a)* |
| yBP2223 | *rnh1Δ rnh201Δ* | *(yBP2213) rnh1::KanMX rnh201::NatMX* | *This study (b)* |
| yBP2216 | *tho (hpr1Δ)* | *(yBP2213) hpr1::HIS3* | *This study (c)* |
| yBP2109 | *tho (mft1Δ)* | *(BY4741) mft1::KanMX* | *Penzo et al<sup>7</sup>* |
| yBP2572 | *rnh201-RED* | *(yBP2213) rnh201-RED-FLAG-HphMX* | *This study (d)* |
| yBP2601 | *mlh3Δ* | *(yBP2213) mlh3::KanMX* | *Euroscarf* |
| yBP2606 | *mlh3Δ mft1Δ* | *(yBP2213) mlh3::KanMX mft1::NatMX* | *This study (e)* |
| yBP2602 | *rad2Δ* | *(yBP2213) rad2::KanMX* | *Euroscarf* |
| yBP2607 | *rad2Δ mft1Δ* | *(yBP2213) rad2::KanMX mft1::NatMX* | *This study (e)* |
| yBP2400 | *Deletion reporter 1n* | *(yBP2213) chrXVI::YPRCTy1-2-URA3-YPRCTy1-4-HphMX* | *This study (f)* |
| yBP2433 | *Deletion reporter 2n* | *chrXVI::YPRCTy1-2-URA3-YPRCTy1-4-HphMX / chrXVI ura3/ura3 his3/his3 leu2/leu2 lys2/LYS2 met15/MET15* | *Mating of BY4742 and yBP2400* |
| yBP2399 | *rnh1Δ rnh201Δ* | *MATalpha ura3 his3 leu2 met15 LYS2 rnh1::KanMX rnh201::NatMX* | *Mating switch in yBP2223* |
| yBP2402 | *Deletion reporter 1n, rnh1Δ rnh201Δ* | *(yBP2400) rnh1::KanMX rnh201::NatMX* | *This study (f)* |
| yBP2435 | *Deletion reporter 2n, rnh1Δ rnh201Δ* | *(yBP2433) rnh1::KanMX/rnh1::KanMX rnh201::NatMX/rnh201::NatMX* | *Mating of yBP2399 and yBP2402* |
| yBP2450 | *Deletion reporter 2n, rnh1Δ* | *(yBP2433) rnh1::KanMX/rnh1::KanMX* | *This study (g)* |
| yBP2451 | *Deletion reporter 2n, rnh201Δ* | *(yBP2433) rnh201::NatMX/rnh201::NatMX* | *This study (g)* |
| yBP2526 | *Deletion reporter 2n, tho (mft1Δ)* | *(yBP2433) mft1::KanMX/ mft1::KanMX* | *This study (g)* |
| yBP2577 | *Deletion reporter 2n, rad2Δ* | *(yBP2433) rad2::KanMX/ rad2::KanMX* | *This study (g)* |
| yBP2578 | *Deletion reporter 2n, rad2Δ rnh201Δ* | *(yBP2433) rad2::KanMX/rad2::KanMX rnh201::NatMX/rnh201::NatMX* | *This study (g)* |
| yBP2573 | *Deletion reporter 2n, mlh3Δ* | *(yBP2433) mlh3::KanMX/mlh3::KanMX* | *This study (g)* |
| yBP2574 | *Deletion reporter 2n, mlh3Δ rnh201Δ* | *(yBP2433) mlh3::KanMX/mlh3::KanMX rnh201::NatMX/rnh201::NatMX* | *This study (g)* |
| yBP2477 | Deletion reporter 2n,<br>*rad52Δ* | (yBP2433) *rad52::KanMX/rad52::KanMX* | This study (g) |
| yBP2479 | Deletion reporter 2n,<br>*rad52Δ rnh201Δ* | (yBP2433) *rad52::KanMX/rad52::KanMX*<br>*rnh201::NatMX/rnh201::NatMX* | This study (g) |
| yBP2579 | Deletion reporter 2n,<br>*rad51Δ* | (yBP2433) *rad51::KanMX/rad51::KanMX* | This study (g) |
| yBP2580 | Deletion reporter 2n,<br>*rad51Δ rnh201Δ* | (yBP2433) *rad51::KanMX/rad51::KanMX*<br>*rnh201::NatMX/rnh201::NatMX* | This study (g) |
| yBP2496 | Deletion reporter 2n,<br>*pol32Δ* | (yBP2433) *pol32::KanMX/pol32::KanMX* | This study (g) |
| yBP2497 | Deletion reporter 2n,<br>*pol32Δ rnh201Δ* | (yBP2433) *pol32::KanMX/pol32::KanMX*<br>*rnh201::NatMX/rnh201::NatMX* | This study (g) |
| yBP2575 | Deletion reporter 2n,<br>*spt3Δ* | (yBP2433) *spt3::KanMX/spt3::KanMX* | This study (g) |
| yBP2576 | Deletion reporter 2n,<br>*spt3Δ rnh201Δ* | (yBP2433) *spt3::KanMX/spt3::KanMX*<br>*rnh201::NatMX/rnh201::NatMX* | This study (g) |
| DLY3211 | *RNH1-HTP* | (BMA64) *RNH1-HTP::KlTRP1* | Aiello et al<sup>8</sup> |
| yBP2458 | *RNH1-HTP hpr1Δ* | (BMA64) *RNH1-HTP::KlTRP1 hpr1::KanMX* | This study (h) |
| ***Candida glabrata*** |  |  |  |
| yBPCG2 | *tho (hpr1Δ)* | *his3 trp1 leu2 hpr1::TRP1* | Bonnet et al<sup>9</sup> |

- A single colony was reisolated from BY4741 as the G<sub>0</sub> parental strain for all subsequent MA lines.
- *RNH1* and *RNH201* complete CDS were deleted sequentially using KanMX and NatMX cassette amplified from pFA6a-KanMX and pFA6a-NatMX, respectively.
- A *hpr1* partial deletion allele marked by the *HIS3* marker was amplified from the original *hpr1* mutant isolate<sup>10,11</sup> and integrated in yBP2213 by homologous recombination.
- A cassette encompassing the *RNH201-RED-FLAG* allele flanked by the HphMX marker was amplified by fusion PCR from YCplac111-rnh201-RED and pFA6a-HphMX and integrated at the *RNH201* locus in yBP2213.
- A *MFT1* deletion cassette was amplified from pFA6a-NatMX and integrated at the *MFT1* locus by homologous recombination within yBP2601 or yBP2602, generating respectively yBP2606 and yBP2607.
- A *URA3* cassette amplified from pRS316 was inserted in-between YPRCTy1-2 and YPRCTy1-4, precisely in the intergenic region between *YPR148c* and *NCE102*. The HphMX cassette (HygroR) was then amplified from pFA6a-HphMX and inserted at the 3' of YPRCTy1-4. Successive integration of the two markers in yBP2213 and yBP2223, respectively, generated yBP2400 and yBP2402.
- Diploid deletion reporter strains were systematically constructed by introducing the mutations of interest into yBP2400 (Deletion Reporter 1n) and the wild-type strain of the opposite mating type (BY4742) via homologous recombination, followed by mating of the resulting strains.
- *HPR1* complete CDS was deleted by homologous recombination in DLY3211 using a cassette amplified from pFA6a-KanMX.

**Suppl. Table 3- Plasmids used in this study**

| Code | Name | Relevant genotype | Source |
| --- | --- | --- | --- |
| pBP7 | *pRS316* | *AmpR, URA3, CEN/ARS* | *Sikorski & Hieter*<sup>12</sup> |
| pBP414 | *pFA6a-KanMX* | *AmpR, KanMX* | *Longtine et al*<sup>13</sup> |
| pBP679 | *pFA6a-NatMX* | *AmpR, NatMX* | *Hentges et al*<sup>14</sup> |
| pBP672 | *pFA6a-HphMX* | *AmpR, HphMX (HygroR)* | *Hentges et al*<sup>14</sup> |
| pBP2272 | *YCplac111-RNH201* | *AmpR, LEU2, CEN/ARS, RNH201-Flag* | *Chon et al*<sup>15</sup> |
| pBP2273 | *YCplac111-rnh201-RED* | *AmpR, LEU2, CEN/ARS, rnh201-P45D Y219A-Flag* | *Chon et al*<sup>15</sup> |

**Suppl. Table 4 - Oligonucleotides used in this study**

| Code | Name | Sequence | Usage |
| --- | --- | --- | --- |
| BP32 | *ACT1-F* | *ACGTTACCCAATTGAACACG* | *Real time*<br>*PCR (RT)* |
| BP33 | *ACT1-R* | *AGAACAGGGTGTTCTTCTGG* |  |
| MZ52 | *Ty1-F* | *ATCGGCAAAAGCACCAAACA* | *Real time*<br>*PCR (RT)* |
| MZ53 | *Ty1-R* | *GTTGTGAACTGGACCAAATATGT* |  |
| MZ127 | *RRP15-F* | *ACGAACAAAGTGATGCTGAAGA* | *Real time*<br>*PCR (RT)* |
| MZ128 | *RRP15-R* | *AGAACCATCATCATGTTTGCTGT* |  |
| MZ131 | *NOC4-F* | *ACGCTTCATACACCTTTCTACAC* | *Real time*<br>*PCR (RT)* |
| MZ132 | *NOC4-R* | *AAAGCTGTTTTCCCGCGG* |  |
| MZ133 | *ASN1-F* | *GCTCCAGATTTGCAAGCCG* | *Real time*<br>*PCR (RT)* |
| MZ134 | *ASN1-R* | *CGTCGTCCAAAGCATCCAAA* |  |
| MZ38 | *Ty1-2-5'-F* | *TATTTTGGTTGCAAAGAAATAAATACCGG* | *Real time*<br>*PCR (DRIP)* |
| MZ39 | *Ty1-2-5'-R* | *AATAATGATGAATTGGGACTTTAAAACCG* |  |
| MZ36 | *Ty1-2-3'-F* | *TTTGAACCTAAAACGCGCCA* | *Real time*<br>*PCR (DRIP)* |
| MZ37 | *Ty1-2-3'-R* | *TCGATTATTTATCCTTCCGTTCATACT* |  |
| MZ42 | *Ty1-4-5'-F* | *TGTCATCATCTTAACACCGTATATGA* | *Real time*<br>*PCR (DRIP)* |
| MZ43 | *Ty1-4-5'-R* | *ACTGACATATCTCATTTTGTGTTGGA* |  |
| MZ40 | *Ty1-4-3'-F* | *TGATACATGTCGCGTTGATATACA* | *Real time*<br>*PCR (DRIP)* |
| MZ41 | *Ty1-4-3'-R* | *ACATTCACCCATTTCTCATTTTGA* |  |
| MZ48 | *intergenicHO-F* | *AGCGACGGTCACATTAGGTT* | *Real time*<br>*PCR (DRIP)* |
| MZ49 | *intergenicHO-R* | *CGCCACAATATACGTACCAGG* |  |
| MZ50 | *YEF3-F* | *ACTCACTCTGCTGAATTCACA* | *Real time*<br>*PCR (DRIP)* |
| MZ51 | *YEF3-R* | *ACCAGCACCTTGACCACTAA* |  |
| RM53 | *RPL15A-F* | *AAGCACCGTGAAGCTAGAGG* |  |
| RM54 | *RPL15A-R* | *TACCAGCCTTGGTGTTGTTG* | Real time<br>PCR (DRIP) |
| AP40 | *LYS2-F* | *GATGGTCACTTGCTGCCTGA* | LYS2<br>amplification<br>for NGS |
| AP41 | *LYS2-R* | *GGGTAGTATGTCACCGACGC* |  |
| - | *Spike-in-F* | *CTATATGCAGCTGGGTGTGTATTTTGTAAACAGAAGTAATTTTCAACTTCT*<br>*AAGCTTTGTATACAAAGCACTGCCGTAGCAATGCCAAGTGCACGTTTCT* | Spike-in<br>(qDRIP) |
| - | *Spike-in-R* | *rArGrArArArCrGrUrGrCrArCrUrUrGrGrCrArUrUrGrCrUrArCrGrGrCrArG*<br>*rUrGrCrUrUrUrGrUrArUrArCrArArArGrCrUrUrArGrArGrUrUrGrArArA*<br>*rArUrUrArCrUrUrCrUrGrUrUrArCrArArArArUrArCrArCrArCrCrArGr*<br>*CrUrGrCrArUrArUrArG* |  |

### Supplementary references

1. Mangione, R. M. *et al.* DNA lesions can frequently precede DNA:RNA hybrid accumulation. *Nat. Commun.* **16**, 2401 (2025).
2. Wahba, L., Costantino, L., Tan, F. J., Zimmer, A. & Koshland, D. S1-DRIP-seq identifies high expression and polyA tracts as major contributors to R-loop formation. *Genes Dev* **30**, 1327–38 (2016).
3. Pelechano, V., Chávez, S. & Pérez-Ortín, J. E. A complete set of nascent transcription rates for yeast genes. *PLoS One* **5**, e15442 (2010).
4. Holstege, F. C. *et al.* Dissecting the regulatory circuitry of a eukaryotic genome. *Cell* **95**, 717–28 (1998).
5. Xu, Z. *et al.* Bidirectional promoters generate pervasive transcription in yeast. *Nature* **457**, 1033–7 (2009).
6. Gomez-Gonzalez, B. & Aguilera, A. Activation-induced cytidine deaminase action is strongly stimulated by mutations of the THO complex. *Proc Natl Acad Sci U S A* **104**, 8409–14 (2007).
7. Penzo, A. *et al.* A R-loop sensing pathway mediates the relocation of transcribed genes to nuclear pore complexes. *Nat. Commun.* **14**, 5606 (2023).
8. Aiello, U. *et al.* Sen1 is a key regulator of transcription-driven conflicts. *Mol. Cell* **82**, 2952–2966.e6 (2022).
9. Bonnet, A. *et al.* Introns Protect Eukaryotic Genomes from Transcription-Associated Genetic Instability. *Mol. Cell* **67**, 608–621.e6 (2017).
10. Aguilera, A. & Klein, H. L. Genetic control of intrachromosomal recombination in *Saccharomyces cerevisiae*. I. Isolation and genetic characterization of hyper-recombination mutations. *Genetics* **119**, 779–790 (1988).
11. Garcia-Rubio, M. *et al.* Different physiological relevance of yeast THO/TREX subunits in gene expression and genome integrity. *Mol Genet Genomics* **279**, 123–32 (2008).
12. Sikorski, R. S. & Hieter, P. A system of shuttle vectors and yeast host strains designed for efficient manipulation of DNA in *Saccharomyces cerevisiae*. *Genetics* **122**, 19–27 (1989).
13. Longtine, M. S. *et al.* Additional modules for versatile and economical PCR-based gene deletion and modification in *Saccharomyces cerevisiae*. *Yeast* **14**, 953–61 (1998).
14. Hentges, P., Van Driessche, B., Tafforeau, L., Vandenhaute, J. & Carr, A. M. Three novel antibiotic marker cassettes for gene disruption and marker switching in *Schizosaccharomyces pombe*. *Yeast* **22**, 1013–9 (2005).
15. Chon, H. *et al.* RNase H2 roles in genome integrity revealed by unlinking its activities. *Nucleic Acids Res.* **41**, 3130–3143 (2013).
